# Strong conservation of spacer lengths in NrdR repressor DNA binding sites

**DOI:** 10.1101/2024.05.27.596032

**Authors:** Saher Shahid, Mateusz Balka, Daniel Lundin, Daniel O. Daley, Britt-Marie Sjöberg, Inna Rozman Grinberg

**Affiliations:** Department of Biochemistry and Biophysics, Stockholm University, SE-10691 Stockholm, Sweden

## Abstract

The ribonucleotide reductase-specific repressor NrdR, from the human pathogens *Listeria monocytogenes* and *Streptococcus pneumoniae*, form tetramers that bind to DNA when loaded with dATP and ATP. If loaded with only ATP they form different oligomeric complexes that cannot bind to DNA. The DNA binding site in *L. monocytogenes* is a pair of NrdR boxes separated by 15-16 bp, whereas in *Streptococcus pneumoniae* the NrdR boxes are separated by 25-26 bp. However, *Streptococcus pneumoniae* NrdR binds stronger to the related *Streptococcus thermophilus* binding sites with NrdR boxes separated by 15-16 bp. This observation triggered a comprehensive binding study of four NrdRs from *L. monocytogenes, Streptococcus pneumoniae, Escherichia coli* and *Streptomyces coelicolor* to a series of synthetic dsDNA fragments where the NrdR boxes were separated by 12-27 bp. All four NrdRs bound well to NrdR boxes separated by 14-17 bp, and also to NrdR boxes separated by 24-27 bp. The worst binding occurred when NrdR boxes were separated by 20 bp. The *in vitro* results were confirmed *in vivo* in *E. coli* for spacer distances 12-27 bp. We conclude that NrdR repressors bind most efficiently when there is an integer number of DNA turns between the center of the two NrdR boxes.

## Introduction

NrdR is a transcriptional repressor protein consisting of a Zn-ribbon domain followed by an ATP-cone (1-3). Its acronym (**n**ucleotide **r**e**d**uctase **r**egulator) stems from the fact that it appears to be specific for the essential enzyme ribonucleotide reductase (RNR) that provides living cells with deoxynucleotide for DNA synthesis and repair (4,5). A balanced supply of deoxynucleotides is a prerequisite for high fidelity DNA synthesis, and the activity of RNRs is controlled at several levels. All RNRs have a specificity regulation that controls the balance between individual dNTP levels (6). A majority of RNRs also have an activity regulation mediated by an ATP-cone, an evolutionarily mobile regulatory domain (7), where binding of ATP activates the enzyme and binding of dATP inhibits enzyme activity. In addition, most bacterial RNRs and a few archeal RNRs are also transcriptionally controlled by NrdR, which also contains an ATP-cone domain (8). In this case, binding of ATP to the ATP-cone of NrdR restricts the repressor from binding to specific NrdR boxes in the promoter regions of RNR operons. With increasing concentrations of dATP, NrdR becomes loaded with both ATP and dATP, which enables binding of NrdR to the NrdR box sequences. The NrdR box motif consists of a pair of 16-bp palindromic sequences separated by a spacer region, where each half of a palindrome binds one Zn-ribbon domain (9). The structure of the functional NrdR tetramer from *Streptomyces coelicolor* loaded with 4 dATP plus 4 ATP molecules and bound to dsDNA was recently solved to high resolution and revealed a constrained DNA helix with sharp kinks at each NrdR box (8).

In this report we have studied the binding of the RNR-specific transcriptional repressor NrdR from two human pathogens: *Streptococcus pneumoniae* and *Listeria monocytogenes*, to NrdR boxes. *Streptococcus pneumoniae* is a major global health burden and a leading cause of death among young children, elderly, and immunocompromised persons (10). *L. monocytogenes* is the third leading cause of death from food-borne illnesses in the United States and a particularly dangerous pathogen among vulnerable groups, such as newborns, pregnant women and the elderly (11). Only two NrdR representatives have been well characterized before: one from *Streptomyces coelicolor* (1,2,8) and the other from *E. coli* (3,12-14). All four bacteria encode two or three different RNR operons, and in three of them (*L. monocytogenes,Streptomyces coelicolor*, and *E. coli*) the spacing between the two NrdR boxes upstream the operons is 15-16 bp, whereas in the *Streptococcus pneumoniae* spacing upstream of both operons is 25-26 bp. This observation raises the question of how similar the mode of RNR regulation by NrdR is in different bacteria. In this study we compare the requirements of NrdRs from these four bacteria from three different phyla. Using a set of synthetic dsDNA fragments with a pair of consensus NrdR boxes with spacers between 12-27 bp we show that all NrdR proteins can bind to motifs with 14-17 bp spacers as well as motifs with 24-27 bp spacers, whereas binding to motifs with other spacer lengths is two orders of magnitude less efficient.

## Materials and Methods

### Plasmids and reagents

The *nrdR* gene from *Listeria monocytogenes* serotype 1/2a str, S10403S (LMRG_RS07775, WP_003723246) and the *nrdR* gene from *Streptococcus pneumoniae* TIGR4 (SP_1713, AAK75791), both cloned in pET30a(+) plasmid using NdeI and XhoI restriction sites, were ordered from GenScript, resulting in pET30a(+)::Lmo*nrdR and pET30a(+)::SpnnrdR*, containing C-terminal hexa-histidine tags. ATP and dATP solutions (100 mM, pH 7.0) were purchased from Thermofisher Scientific. c-diAMP was obtained from Jena Bioscience, Germany in lyophilized form and resuspended in water to 10 mM solution. ADP (Sigma Aldrich) powder was dissolved in water to get 100 mM solution and pH was adjusted to 7.0.

### Protein expression and purification

*E. coli* BL21(DE3) were transformed with the pET30a(+)::Lmo*nrdR* or *pET30a(+)::SpnnrdR*. Single colony of the transformed cells was used to inoculate the LB medium supplemented with kanamycin (50 μg/ml) and grown overnight at 37 °C. This overnight culture was used to inoculate LB medium to OD_600_ 0.1 and the cells were allowed to grow under vigorous shaking at 37 °C. When OD_600_ of 0.8-0.9 was achieved, the culture media was cooled down on ice and 0.5 mM IPTG and 0.1 mM Zn(CH_3_CO_2_)_2_ were added. The cultures were grown overnight at 20 °C and the cells expressing LmoNrdR or SpnNrdR were harvested by centrifugation.

Cell pellet was resuspended in lysis buffer containing 1M NaCl, 20% glycerol, 10 mM Imidazole, 2 mM DTT and 0.05 M Tris-Cl pH 8.5 at 4°C for LmoNrdR and 0.05 M HEPES pH 7.2 for SpnNrdR. Phenylmethylsulfonyl fluoride (PMSF) was added to 1 mM concentration to the cell suspension, which was then sonicated in an ultrasonic processor (Misonics) until a clear lysate was obtained. The lysate was centrifuged at 18,000 x g at 4°C for 45 min and the supernatant was used for protein purification.

For purification, the supernatant was loaded into HisTrap FF Ni-Sepharose column (Cytiva) which was pre-equilibrated with lysis buffer (without PMSF). After loading the protein, the column was extensively washed with buffer containing 10 mM and 60 mM imidazole. The his-tagged LmoNrdR/SpnNrdR were eluted with elution buffer containing 0.5 M NaCl, 20% glycerol, 0.5 M Imidazole, 2 mM DTT and 0.05 M Tris-Cl pH 8.5 at 4°C for LmoNrdR and 0.05 M HEPES pH 7.2 for SpnNrdR. The purified protein was then desalted using HiPrep 26/10 Desalting column (Cytiva) equilibrated with storage buffer (0.05 M Tris-Cl (pH 8.5 at 4°C), 0.5 M NaCl, 20% glycerol, 1 mM TCEP for LmoNrdR and 0.05 M HEPES (pH 7.2 at RT), 0.5 M NaCl, 20% glycerol,1 mM TCEP for SpnNrdR) and stored at -80°C. Protein concentration was determined using Coomassie Plus protein assay reagent (ThermoFisher Scientific) using a BSA standard curve.

To remove traces of bound nucleotides from LmoNrdR, the purified protein was subjected to hydrophobic interaction chromatography (HIC) as described previously (8) using HiTrap Phenyl HP column (Cytiva) in buffer (0.025 M Bis Tris Propane (pH 6.5), 0.75 M ammonium sulfate and 0.1 mM TCEP). After applying the protein, the column was washed extensively with the same buffer (50 CV) and the protein was eluted in the same buffer without ammonium sulfate. The eluted protein was then desalted using a PD10 column (Cytiva) in buffer containing 0.025 M Bis Tris Propane pH 6.5, 0.3 M NaCl, 10% glycerol and 1 mM TCEP). Protein concentration and recovery after HIC was 0.8 mg/ml and ∼20%, respectively. The apo-LmoNrdR was aliquoted and stored in -80 °C until used.

### Microscale thermophoresis

Microscale thermophoresis (MST) was used to study the binding of NrdR proteins to strain specific and synthetic NrdR boxes. Oligonucleotides of 57-69 bp containing two NrdR boxes, the spacer region between them and flanking regions of 5 bp were ordered from Merck (Germany). The 5’ end of sense strand of each oligonucleotide was labeled with Cyanine 5 dye (Cy5) by the manufacturer, while the antisense strand was unlabeled. Oligonucleotides were either obtained as 100 µM solution in Tris-EDTA buffer or in lyophilized form and resuspended in 50 mM Tris-Cl (pH 8.0), 50 mM NaCl and 1 mM EDTA. Single stranded oligonucleotides were annealed by mixing 50 and 57.5 pmoles of labeled and unlabeled oligonucleotides respectively, in 50 ul buffer (10 mM Tris-Cl pH 8.0 and 50 mM NaCl) using a thermoblock. Annealing program involved incubation for 5 min at 95°C followed by gradual cooling to 25°C using 140 cycles of - 0.5°C and 45 sec per cycle, resulting in 1 µM double stranded DNA. Integrity of the annealed oligonucleotides was determined by application to Mini-PROTEAN native 5% polyacrylamide TBE gel (Biorad), staining with SYBR safe DNA gel stain (Thermo Fischer Scientific) and analysis using gel imager system C600 (Azure Biosciences) at wavelengths suitable for detection of Cy5 and Cy3 emission signals (Supplementary Figure 1). Oligonucleotides used to assay NrdR binding to native *L. monocytogenes, Streptococcus pneumoniae* and *Streptococcus thermophilus* promoters are listed in supplementary table 1. The synthetic (Synt) oligonucleotides with different spacer lengths are listed in supplementary tables 2 and 3 (for sense and antisense oligonucleotides respectively).

MST for analyzing the binding of LmoNrdR and SpnNrdR to strain-specific NrdR boxes and synthetic oligonucleotides containing NrdR boxes with spacers of different lengths was performed using Monolith NT.115 instrument (Nanotemper Technologies, Germany), while the binding of ScoNrdR and EcoNrdR to the synthetic oligonucleotides was performed using Monolith NT. Automated instrument (Nanotemper Technologies, Germany).

For LmoNrdR, MST buffer contained 0.025 M Bis Tris Propane (pH 6.5), 5% glycerol, 100 mM KCl, 5 mM MgCl_2_, 1 mM DTT and 0.05% Tween 20. For assaying binding of LmoNrdR to native NrdR boxes in the presence of different nucleotide effectors, apo-LmoNrdR, obtained by HIC, was supplemented with the nucleotides of choice. For assaying binding to synthetic oligonucleotides, protein purified by nickel affinity chromatography, supplemented with dATP and ATP was used. Either one nucleotide at 1 mM concentration or two nucleotides at 0.5 mM concentration each were included in MST buffer. To study the binding of LmoNrdR to oligonucleotides containing native NrdR boxes, medium MST power, 90% of excitation power and 2 nM DNA was used. For the analysis of LmoNrdR binding to synthetic oligonucleotides with different spacer lengths, high MST, 95% excitation power and 0.5 nM DNA were used.

For SpnNrdR the buffer was 0.025 M HEPES (pH 7.2), 250 mM NaCl, 15% glycerol, 2.5 mM MgCl_2_, 2 mM DTT, 0.1 mg/ml BSA, 0.05% Tween 20, 0.5 mM ATP and/or 0.5 mM dATP. MST was done at medium MST power, 90% excitation power and with 5 nM DNA.

For EcoNrdR the buffer was 0.025 M Tris-Cl (pH 8.5 at 4 °C), 100 mM NaCl, 7 mM MgCl_2_, 1mM DTT, 0.025% Tween20, 1 mM ATP and 1 mM dATP. MST was done at high MST power with 50% excitation power using 10 nM DNA.

For ScoNrdR, the buffer was 0.025 M Tris-Cl (pH 8.0), 150 mM NaCl, 5% glycerol, 10 mM MgCl_2_, 2 mM DTT, 0.05% Tween 20, 1 mM ATP and 1 mM dATP. MST was done at medium MST power, 40% excitation power and with 10 nM DNA.

The results were analyzed using the MO.Affinity Analysis v2.3 software (Nanotemper) with default parameters. K_D_ and standard deviation were calculated using fits from at least three individual titrations. In individual cases the K_D_s couldn’t be reliably determined since the titration curves didn’t reach a plateau or no binding was observed (see titration curves in supplementary materials). The indicated K_D_s, derived from these curves, are therefore only estimates. These titrations were performed at least twice for each of the tested promoters.

### Analytical size exclusion chromatography

In order to see the oligomerization states of the protein in the presence or absence of different nucleotides, size exclusion chromatography (SEC) was done at 4 °C using Superdex S200 10/300 Increase or Superdex S200 3.2/300 Increase (Cytiva) attached to an Äkta prime system (Cytiva). Molecular weights were estimated from the calibration curve which was derived from the globular protein standards using both the high and low molecular weight SEC marker kits (Cytiva). Standard deviations were calculated using data from at least three runs.

For LmoNrdR protein, nucleotides/combinations of nucleotides added to the protein included dATP + ATP, ATP, ADP, dATP + ADP and c-di-AMP. The column was equilibrated with 25 mm BTP (pH 6.5), 5% glycerol, 5 mM MgCl_2_, 100 mM KCl and 1 mM DTT. One (0.2 mM) or combination of two nucleotides (0.1 mM each) (ATP, dATP, ADP and c-di-AMP) were included in the buffer. The protein (50 µM) was incubated for 10 min at room temperature with 1 mM nucleotide and 5 mM MgCl, centrifuged and 100 µl sample was injected to the column. Runs of apo-NrdR were performed without addition of nucleotides to protein or buffer.

For SpnNrdR protein, the column was equilibrated with 25mM HEPES (pH 7.2), 2.5mM MgCl2, 15% glycerol, 250 mM NaCl and 0.1 mM TCEP and either ATP and dATP 0.1 mM each or 1 mM ATP. SpnNrdR protein (50 µM) was incubated for 10 min at room temperature with either 0.5 mM ATP, 0.5 mM dATP and 2.5 mM MgCl or 1 mM ATP and 2.5 mM MgCl, centrifuged and 100 µl sample was injected to the column. Runs of apo-NrdR were performed without addition of nucleotides to protein or buffer.

### Promoter activity assay for *in vivo* analysis of repressor binding

Double-stranded DNA fragments (201-216 bp) containing the ATG start codon of *E. coli* K-12 strain MG1655 *nrdHEF* gene and 143 bases upstream, i.e. containing the NrdR and Fur binding sites, (P_nrdHEF_) were ordered from GenTitan™ Gene Fragments Service (Genscript). The sequence of the region containing the NrdR boxes and the spacer between them in each of the constructs are given in supplementary table 4. The fragments were cloned into the pSEVA631(Sp) [P_ibpA_-gfp_ASV_] plasmid (15) using the *in vivo* cloning method (16). In short, the pSEVA plasmid was amplified by PCR using the Q5 High-Fidelity DNA Polymerase (New England Biolabs) and the Fwd_GFP_rep_AN (ATGCGTAAAGGAGAAGAACTTTTCAC) and Rev_GFP_rep_AN primers (TTAATTAAAGGCATCAAATAAAACGAAAGG). This PCR removed the region encoding P_ibpA_. The PCR product was digested with DpnI, separated by agarose gel electrophoresis, and purified using DNA gel extraction kit (GeneJET, Thermo Scientific), mixed briefly at room temperature with P_nrdHEF_ dsDNA fragments (one at a time) and transformed into the chemically competent MC1061 strain (str.K-12F^−^ λ^−^ Δ*(ara-leu)7697* [*araD139*]B/rΔ*(codB-lacI)3 galK16 galE15* e14^−^ *mcrA0 relA1 rpsL150*(Str^R^) *spoT1 mcrB1 hsdR2*(*r*^−^*m*^+^)). The integrity of the resulting plasmids pSEVA631(Sp) [P_nrdHEF_-gfp_ASV_] was verified by sequencing carried out by Eurofins Genomics (Germany).

A single colony of MC1060 harboring pSEVA631(Sp) [P_nrdHEF_-gfp_ASV_] was used to inoculate 500 µL of LB media (Miller) containing 50 µg/mL spectinomycin. Cultures were grown at 37°C with shaking at 185 rpm for 16 to 20 hours. A 50 µL aliquot was back-diluted into 5 mL of fresh LB containing 50 µg/mL spectinomycin in a 5 mL 24-well plate. Cultures were grown at 37°C with shaking at 185 rpm to an OD_600_ of approximately 0.8. Aliquots of 2 - 5 mL bacterial culture were collected by centrifugation at 3220 × g for 15 min. The culture medium was removed, and the pelleted cells were resuspended in 200 μL of buffer (50mM Tris–HCl pH 8.0, 200mM NaCl, 15mM EDTA). The cell suspension was transferred to a 96-well optical bottom black-wall plate (Thermo Scientific) and the fluorescence was determined in a SpectraMax Gemini EM plate reader (Molecular Devices, U.K.). Fluorescence from pSEVA631(Sp) [P_nrdHEF_-gfp_ASV_] was measured with an excitation wavelength of 475 nm and emission wavelength of 515 nm. A long-pass emission cut-off filter of 495 nm was used to reduce background. Fluorescence values were normalized by the optical density and the volume of the sample (OD_600_). These values were obtained from a 200 µL aliquot of bacterial culture using a SpectraMax *m2e* plate reader (Molecular Devices, U.K.) at 600 nm.

### Bioinformatics

After identifying potential RNR operons – genes annotated as RNRs by Prokka (v. 1.14.6, (17)), separated by no more than 300 nucleotides – in all species representative bacterial genomes in the GTDB (release 08-RS214, (18)) 500 nucleotides upstream of each operon were extracted. Each fragment was searched with a regular expression for a pair of NrdR boxes separated by a spacer of no more than 50 nucleotides (“C.[AT].[AT][AT].[AT].G.{1,50}?C.[AT].[AT][AT].[AT].G”) and the pair closest to the start of the first coding region was selected. Spacers with a length between 20 and 23 (corresponding to a box distance of 14-17 bp) were considered “short” and those between 30 and 33 (corresponding to a box distance of 24-27 bp) were considered “long”. Finally, each class of lengths was counted.

## Results

### Binding of *Streptococcus pneumoniae* and *L. monocytogenes* NrdRs to their respective RNR promoter regions

The human pathogen *Streptococcus pneumoniae* (19,20) encodes two RNR operons, *nrdHEF* coding for an aerobic class Ib RNR and *nrdDG* coding for an anaerobic class III RNR. Interestingly, in the promoter region of *nrdDG* the predicted NrdR boxes are separated by a spacer region of 25 bp, i.e. approximately one helix turn longer than the common spacer region of 15 to 16 bp (Fig. 1). Box 2 includes the -35 sequence of the RNA polymerase binding site, while the -10 region is downstream of box 2. In the promoter region of the *nrdHEF* operon we find three NrdR boxes. Boxes 0 and 1 are separated by 24 bp and boxes 1 and 2 by 26 bp (Fig. 1). Both the -35 and -10 sequences are downstream of box 2.

**Figure 1.**
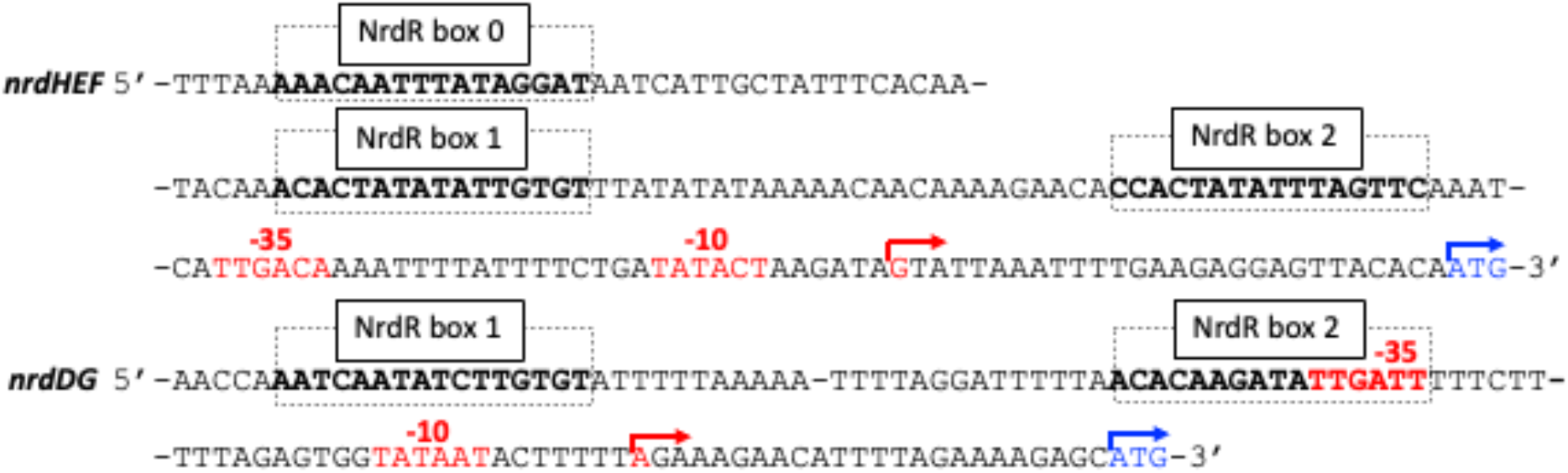
Promoter regions of *nrdHEF* and *nrdDG* in *Streptococcus pneumoniae*. The boldface black/red and boxed regions refer to NrdR boxes 0, 1, and 2 of the *nrdHEF* operon, and NrdR boxes 1 and 2 of the *nrdDG* operon. Red letters and arrows indicate the predicted -35 and - 10 regions and the transcription start sites, and blue letters and arrows show translation start sites (21). The ATG start codon is 68 bp downstream of Box 2 in the *nrdHEF* promoter region and 53 bp downstream of Box 2 in the *nrdDG* promoter region.

The human pathogen *L. monocytogenes* (22,23) encodes two RNR operons, *nrdABI-trxL* (here called *nrdAB)*, coding for an aerobic class I RNR, and *nrdDG*, coding for an anaerobic class III RNR (24). Upstream of the *nrdDG* operon is a pair of NrdR boxes that overlaps the RNA polymerase binding site and the NrdR boxes are separated by a 16 bp spacer sequence (Fig. 2). The upstream region of the *nrdAB* operon instead includes four NrdR boxes, one proximal pair and one distal pair (Fig. 2). The promoter-distal pair of NrdR boxes are separated by a 15 bp spacer sequence and the promoter-proximal pair of boxes are separated by a 16 bp spacer sequence. The proximal NrdR box motif overlaps with the RNA polymerase binding site.

**Figure 2.**
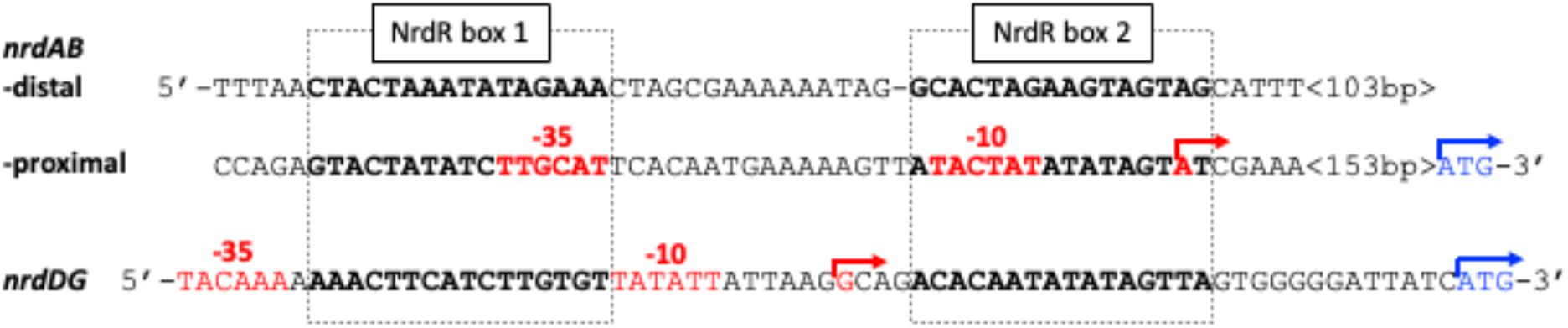
Promoter regions of *nrdABI-trxL* and *nrdDG* in *L. monocytogenes*. The boldface black/red and boxed regions refer to NrdR boxes 1 and 2. Red letters and red arrows indicate the predicted -35 and -10 regions and the transcription start sites (25,26); blue letters and arrows show translation start sites. The ATG start codon is 157 bp downstream of proximal Box 2 in the *nrdAB* promoter region and 13 bp downstream of Box 2 in the *nrdDG* promoter region.

*Streptococcus pneumoniae* NrdR (SpnNrdR) loaded with dATP and ATP binds comparatively weakly to dsDNA fragments corresponding to the long NrdR box motifs upstream of the *nrdDG* operon and the *nrdHEF* (box 1 and 2) operon from *Streptococcus pneumoniae* (Table 1, Supplementary fig. 2a). In addition, dATP-plus ATP-loaded SpnNrdR binds even less well (K_D_ >7 times higher) to the NrdR boxes 0 and 1 upstream of the *nrdHEF* promoter region.

**Table 1.** Effects of nucleotides on binding of SpnNrdR to RNR promoter regions from *Streptococcus pneumoniae*.

| **Nucleotide(s)** | ***nrdHEF* Box 0+1**<br>**K<sub>D</sub> (μM)** | ***nrdHEF* Box 1+2**<br>**K<sub>D</sub> (μM)** | ***nrdDG***<br>**K<sub>D</sub> (μM)** |
| --- | --- | --- | --- |
| dATP + ATP | >69 | 11 ± 0.9 | 15 ± 3 |
| ATP | No binding | >91 | >3000 |

Interestingly, dATP-plus ATP-loaded SpnNrdR binds approximately 10 times stronger to shorter NrdR box motifs, with a spacer distance of 16 bp, upstream of the *nrdHEF* and *nrdDG* operons in *Streptococcus thermophilus* (Table 2, Supplementary fig. 2b). As expected, ATP-loaded SpnNrdR has extremely low affinity to all dsDNA fragments tested (Table 1, Table 2, Supplementary fig. 2).

**Table 2.** Binding of SpnNrdR in presence of dATP+ATP to RNR promoter regions from *Streptococcus thermophilus*.

| **NrdR binding site** | **SpnNrdR + dATP + ATP**<br>**K<sub>D</sub> (μM)** | **SpnNrdR + ATP**<br>**K<sub>D</sub> (μM)** |
| --- | --- | --- |
| *S. thermophilus nrdHEF* NrdR boxes | 0.80 ± 0.25 | >42 |
| *S. thermophilus nrdDG* NrdR boxes | 1.15 ± 0.32 | >80 |

*L. monocytogenes* NrdR (LmoNrdR) loaded with dATP and ATP can bind to dsDNA fragments from all three sets of NrdR boxes with extremely high affinity (low nM range). Addition of other nucleotides resulted in significantly lower binding affinities or no binding at all (Table 3, Supplementary fig. 3). The secondary metabolite di-cyclic-AMP (c-di-AMP) was earlier reported to bind LmoNrdR (27), but neither c-di-AMP or its combination with ATP or dATP promoted binding of NrdR to DNA in our experiments (Table 3, Supplementary fig. 3). Similarly, *Streptomyces coelicolor* NrdR (ScoNrdR) has the ability to bind c-di-AMP with a K_D_ ∼3 µM but has low affinity to DNA when loaded with c-di-AMP (8).

**Table 3.**
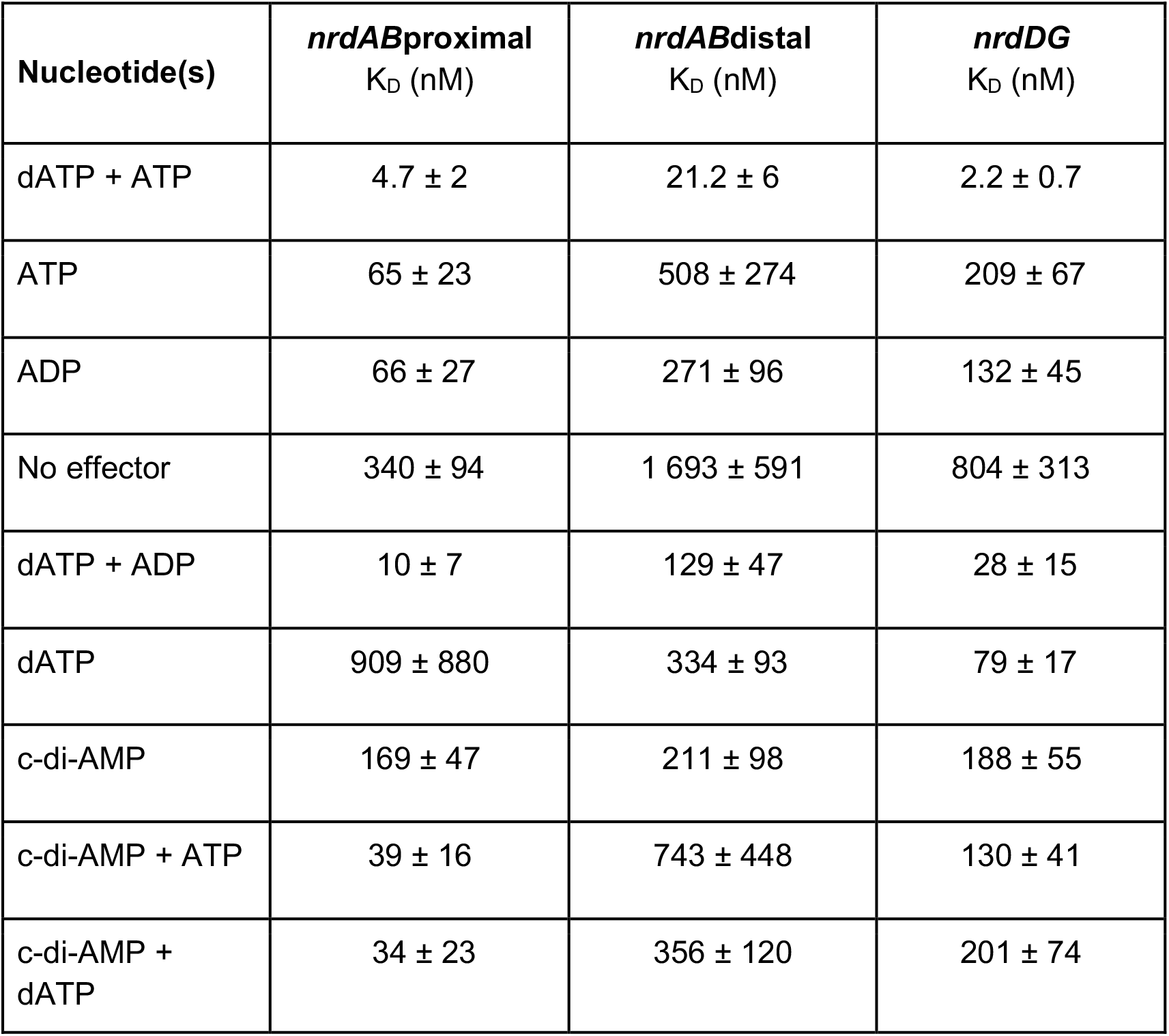
Effects of nucleotides on binding of LmoNrdR to RNR promoter regions from *L. monocytogenes*.

We performed size exclusion chromatography (SEC) to determine the oligomeric state of SpnNrdR and LmoNrdR in the presence of different nucleotides. SpnNrdR is an equilibrium of dimers and tetramers in its apo form (in the absence of nucleotides), a tetramer in presence of a combination of dATP and ATP, and primarily a high molecular weight oligomer in the presence of ATP (Fig. 3a, Table 4). LmoNrdR eluted as a dimer in its apo form, and as a tetramer in presence of dATP and ATP, and as an octamer in presence of ATP (Fig. 3b, Table 4). LmoNrdR also eluted as a tetramer in the presence of ADP, a combination of dATP+ADP, or c-di-AMP (Supplementary fig. 4). These results are in line with the previously reported oligomeric states for *Streptomyces coelicolor* and *E. coli* NrdR (8,28).

**Figure 3.**
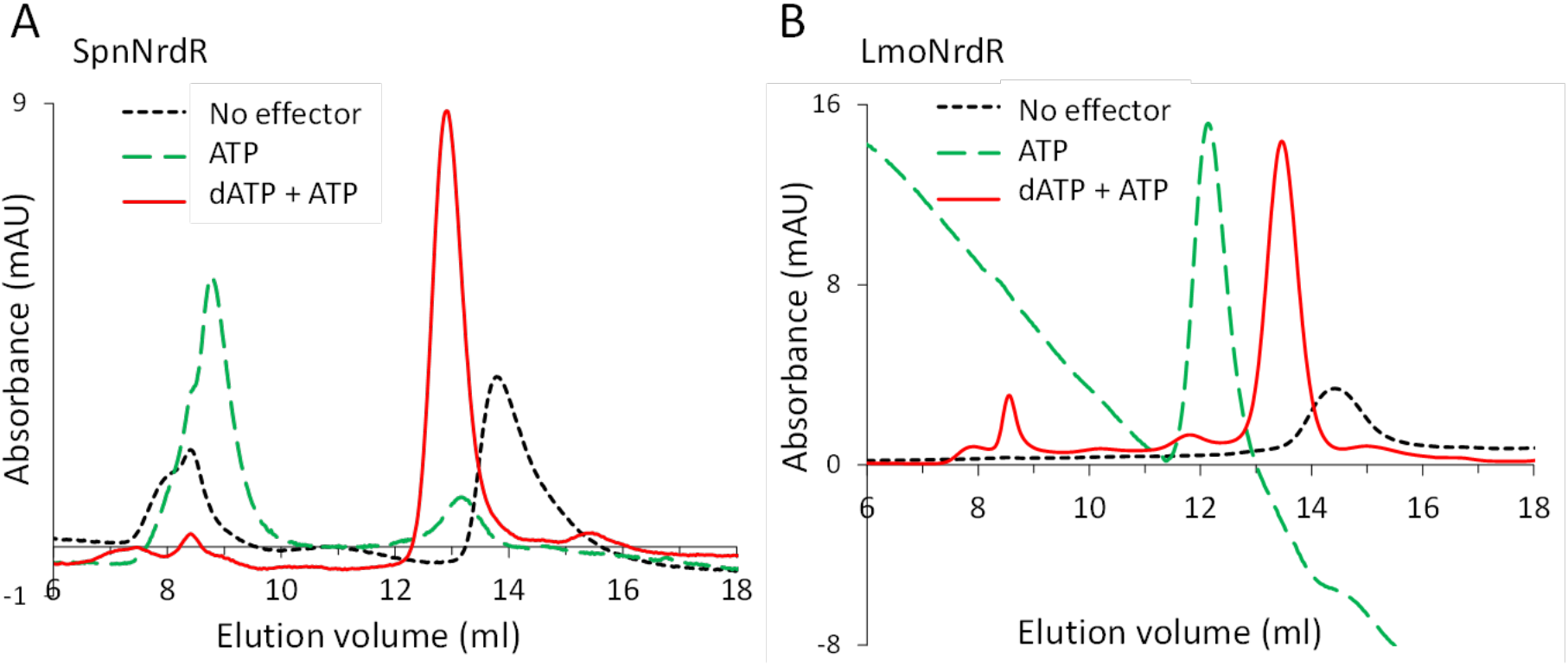
Size exclusion chromatography in presence of different nucleotide combinations. Representative chromatograms of a) SpnNrdR and b) LmoNrdR performed in the presence of ATP (dashed green), dATP + ATP (solid red) and no nucleotides added (dotted black).

**Table 4.**
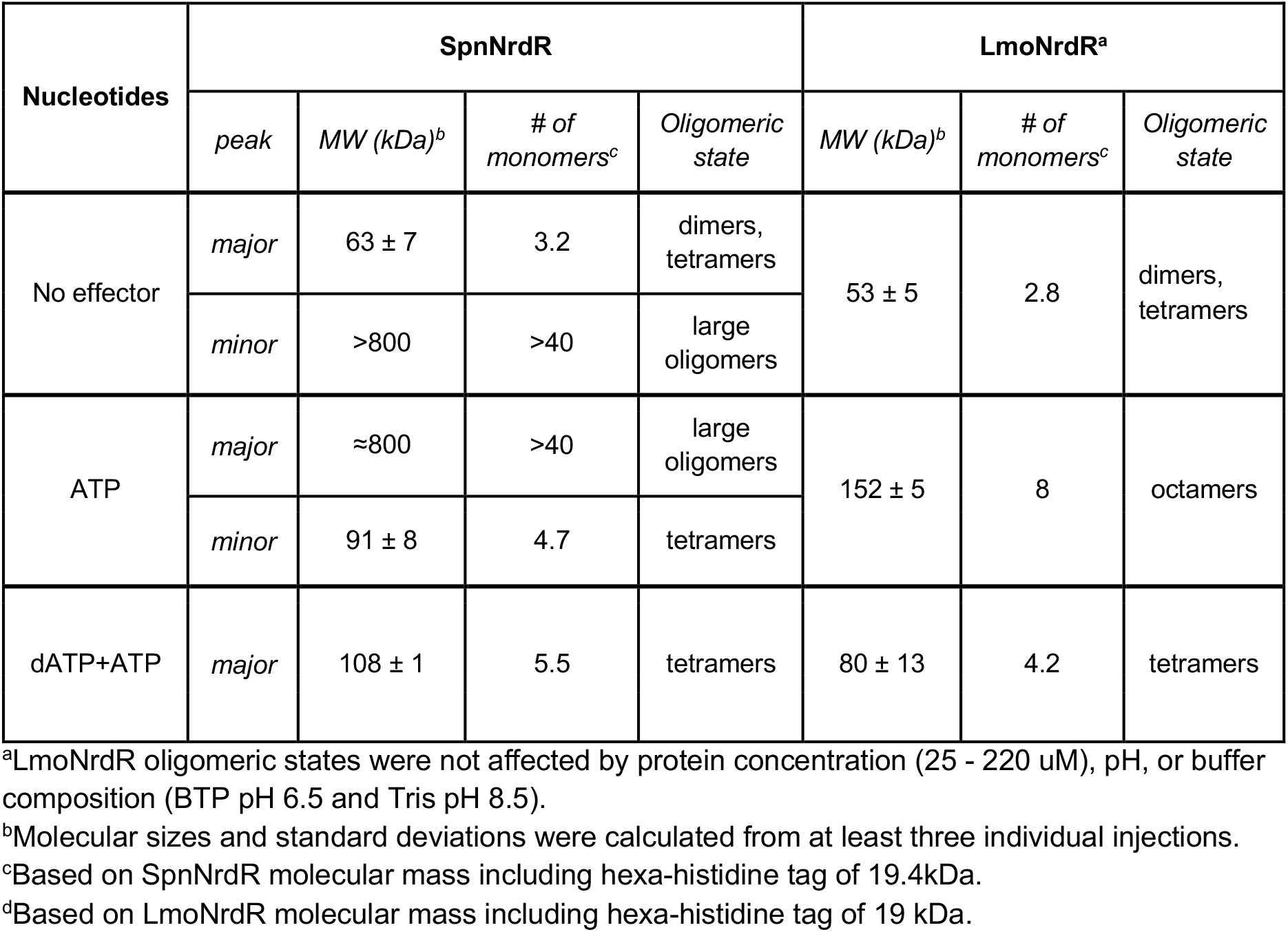
Summary of oligomeric states of SpnNrdR and LmoNrdR in presence of different effectors.

### Spacer lengths between pairs of NrdR boxes in upstream regions of RNR operons in *Streptococcus* spp and *Listeria* spp

A bioinformatic analysis of NrdR box motifs in RNR promoter regions in *Streptococcus* spp and *Listeria* spp revealed that a majority of *Streptococci* have long spacers between the NrdR boxes whereas a majority of *Listeria* have short spacers (Table 5).

**Table 5.** Fractions of long and short spacer lengths in NrdR binding sites upstream RNR operons in *Streptococcus* spp and *Listeria* spp. Long spacers are 24-27 bp and short spacers are 14-17 bp.

| Genus | Operon | Total NrdR box pairs | Long spacers | Short spacers | Box pairs with short spacers (%) |
| --- | --- | --- | --- | --- | --- |
| *Streptococcus* | *nrdD* | 402 | 282 | 59 | 17 |
|  | *nrdHEF* | 364 | 252 | 70 | 19 |
| *Listeria* | *nrdD* | 22 | 4 | 16 | 73 |
|  | *nrdAB* | 21 | 3 | 17 | 81 |

The vast majority of *Streptococcus* species had comparable spacer lengths in both promoters i.e. species with long spacer between boxes in *nrdHEF* promoter also had a long spacer in *nrdDG* promoter and those with a short spacer in *nrdHEF* promoter had a short spacer in *nrdDG* promoter (Fig. 4). There were, however, some exceptions e.g. *S. suis*, with a short spacer in *nrdEF* promoter and long spacer in *nrdDG* promoter. NrdR boxes in *nrdDG* promoter and boxes 1 and 2 in *nrdEF* promoter are conserved in *Streptococcus spp*. (Fig. 4), while box 0 in *nrdEF* promoter is not conserved (data not shown). SpnNrdR binds very weakly to *nrdHEF* box 0 and 1 suggesting that it doesn’t recognize this pair of boxes *in vivo*.

**Figure 4.**
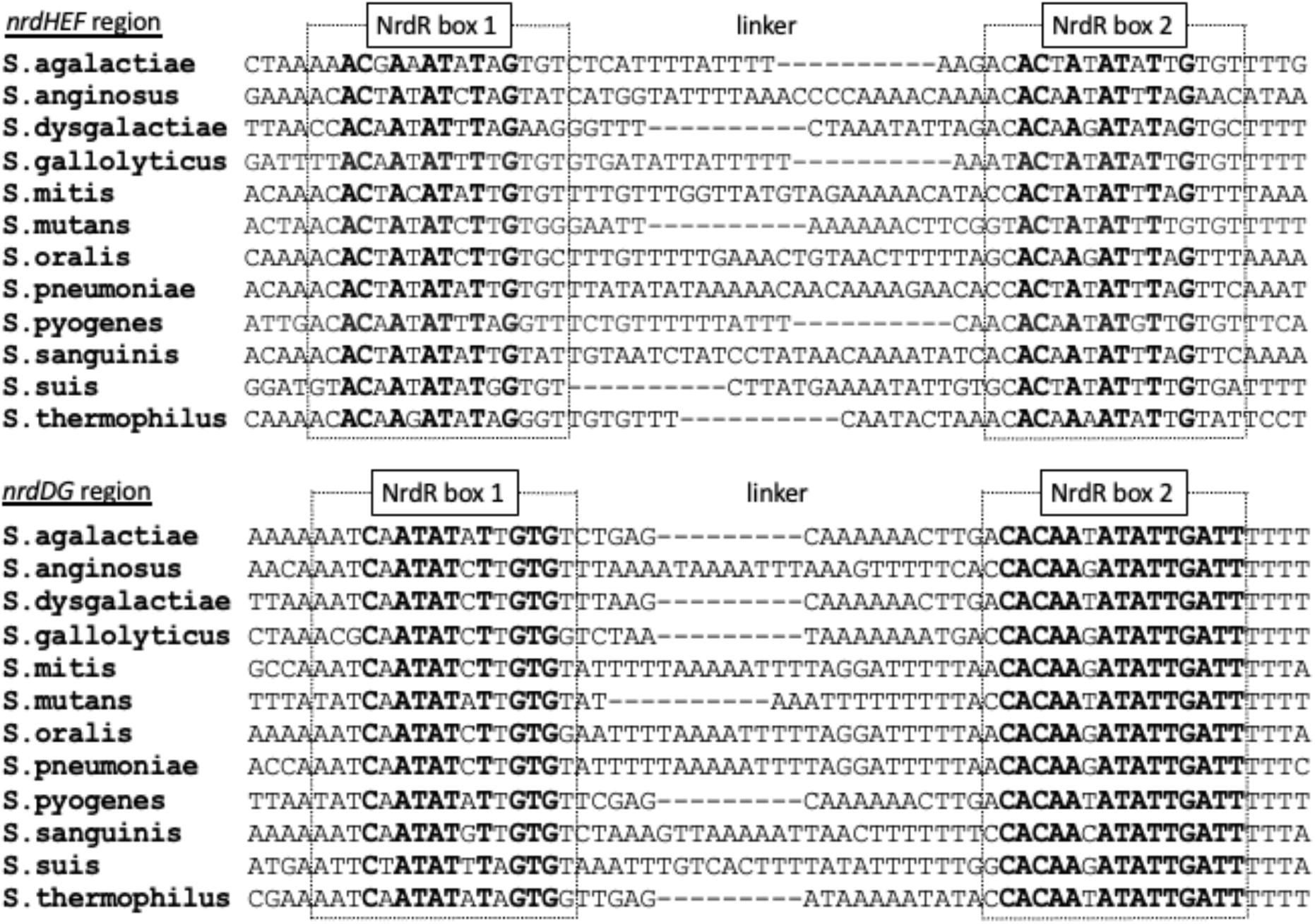
NrdR boxes upstream of *nrdDG* genes and *nrdHEF* genes in selected *Streptococci*. Conserved positions in the box regions are in boldface.

### Effect of spacer region length between the NrdR boxes on NrdR binding affinity

In the *L. monocytogenes nrdAB* promoter region two sets of RNR boxes are found. In the distal boxes, the spacer between two boxes is 15 nucleotides, while in the proximal boxes and in the *nrdDG* promoter region the distance is 16 nucleotides. Our binding results show that LmoNrdR has 5-10 times stronger binding to the *nrdDG* binding site and the *nrdAB* proximal bindings site than it has to the *nrdAB* distal site (Table 1). In the *Streptococcus pneumoniae nrdDG* and *nrdHEF* promoters the spacer lengths between NrdR boxes are 25-26 bp, whereas the related *Streptococcus thermophilus nrdDG* and *nrdHEF* promoters have a spacer length of 16 bp, and paradoxically bind SpnNrdR approximately 10 times stronger than do the homologous binding sites in *Streptococcus pneumoniae* (Tables 2 and 3). We therefore decided to explore the effect of the spacer length on NrdR binding affinity using synthetic NrdR boxes separated by a synthetic spacer region (12 to 27 bp in length; Supplementary tables 2-3). The results demonstrate that LmoNrdR is bound tightly to NrdR-boxes with a spacer region between 14-18 bp with K_D_s between 1 nM and 3 nM but bound significantly weaker to shorter or longer spacers (K_D_s between 11 nM and 122 nM) (Fig. 5A, Table 6, Supplementary fig. 5). Interestingly, SpnNrdR appeared to have a sharper spacer length requirement with K_D_s between 41 nM and 55 nM for spacers of 15-16 bp (Fig. 5B, Table 6, Supplementary fig. 6).

**Figure 5.**
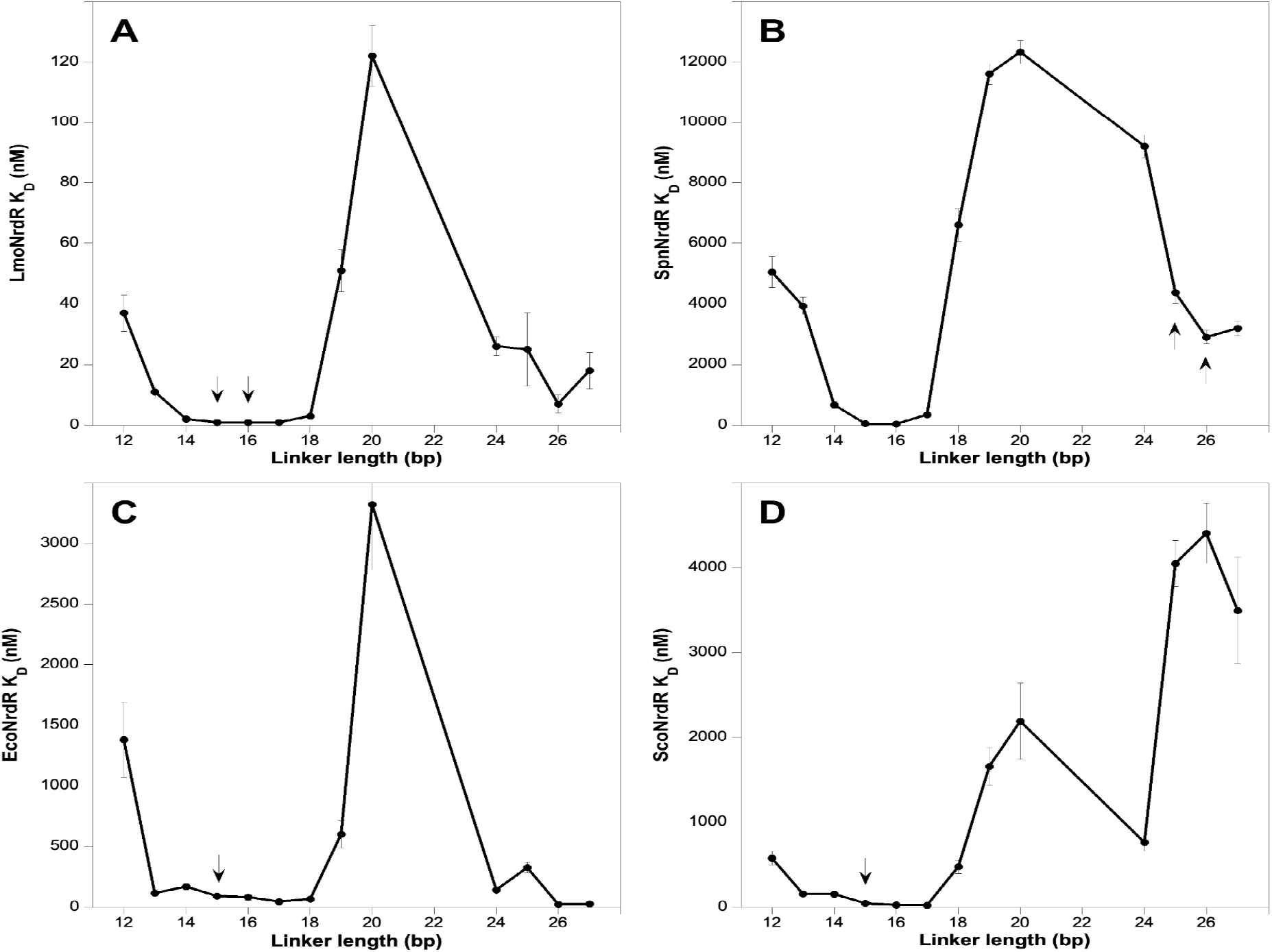
Effect of spacer length in synthetic dsDNA NrdR box pairs on the binding constant of LmoNrdR (A), SpnNrdR (B), EcoNrdR (C), and ScoNrdR (D). Arrows denote the spacer lengths of the binding sites in the native RNR upstream regions in each case.

**Table 6.** Binding affinities of NrdRs from *L. monocytogenes, Streptococcus pneumonia, E. coli* and *Streptomyces coelicolor* to synthetic NrdR boxes having different spacer lengths between the boxes.

| Spacer region<br>(bp) | K<sub>D</sub> (nM) |  |  |  |
| --- | --- | --- | --- | --- |
|  | LmoNrdR | SpnNrdR | EcoNrdR | ScoNrdR |
| 12 | 37 ± 6 | 5 057 ± 511 | 1 381 ± 309 | 580 ± 79 |
| 13 | 11 ± 1.8 | 3 936 ± 290 | 117 ± 24 | 158 ± 19 |
| 14 | 2 ± 0.5 | 699 ± 109 | 169 ± 27 | 154 ± 28 |
| 15 | 1 ± 0.2 | 55 ± 14 | 91 ± 14 | 45 ± 13 |
| 16 | 1 ± 0.4 | 41 ± 11 | 82 ± 22 | 28 ± 7 |
| 17 | 1 ± 0.4 | 340 ± 83 | 45 ± 6 | 21 ± 7 |
| 18 | 3 ± 0.6 | 6 603 ± 536 | 66 ± 12 | 476 ± 80 |
| 19 | 51 ± 7 | 11 608 ± 340 | 601 ± 113 | 1 658 ± 221 |
| 20 | 122 ± 10 | 12 330 ± 388 | 3 322 ± 540 | 2 191 ± 449 |
| 24 | 26 ± 3 | 9 200 ± 374 | 141 ± 34 | 763 ± 104 |
| 25 | 25 ± 12 | 4 371 ± 343 | 326 ± 44 | 4 051 ± 272 |
| 26 | 7 ± 3 | 2 906 ± 225 | 24 ± 7 | 4 404 ± 355 |
| 27 | 18 ± 6 | 3 194 ± 238 | 27 ± 8 | 3 496 ± 626 |

To find out if this relaxed dependence on spacer length is specific for LmoNrdR and SpnNrdR or a general feature of other NrdRs we also tested binding of the NrdR proteins from *E. coli* (EcoNrdR) and *Streptomyces coelicolor* (ScoNrdR) to the set of synthetic NrdR boxes with varying spacer lengths. These four NrdR proteins have 39-52% sequence identity (Supplementary fig. 7). For EcoNrdR, good binding was observed for spacers of 15-18 bp with K_D_s between 45 and 91 nM (Fig. 5C, Supplementary fig. 8), and for ScoNrdR best binding was obtained for spacers of 15-17 bp with K_D_s between 21 to 45 nM (Fig. 5D, Supplementary fig. 9).

A spacer length of 15-16 bp would result in approximately one and a half turns of the DNA helix between the NrdR boxes (i.e. approximately three turns of the DNA helix between the center of the two NrdR boxes in the binding site). To study whether the different NrdR proteins would also accept a promoter region with an extra DNA helix turn between the center of the NrdR boxes, we tested binding to synthetic fragments with spacer region lengths between 24 bp and 27 bp. It is obvious from table 6 and figure 5 that EcoNrdR can bind to the DNA fragments with 26 and 27 bp spacer with two-fold higher affinity than to fragments with a 17 bp spacer. LmoNrdR and ScoNrdR were less tolerant and had K_D_s for the 26 and 24 bp long spacers that were 7 and 36 times worse than the best K_D_s that were observed to fragments with optimal short spacers. SpnNrdR turned out to be the least tolerant of the tested NrdRs with a 70-fold higher K_D_s for the 26 bp long DNA fragment compared to its K_D_ for the DNA fragment with a 16 bp spacer.

### *In vivo* confirmation of spacer length results

To assess the relevance of the *in vitro* results of different spacer lengths we designed a set of experiments in *E. coli* cells. Reporter plasmids with the coding sequence for the green fluorescent protein (GFP) were linked to the upstream region of the *E. coli nrdEF* operon. During exponential growth the *nrdEF* operon is strongly repressed by the transcription factors NrdR and Fur, and it is derepressed at stationary growth and/or iron starvation (3,29). The expression of GFP will therefore be low in our reporter system if NrdR can bind to the NrdR boxes. Expression will be higher if NrdR is prevented from binding by an unfavorable spacing between the NrdR boxes or if the NrdR boxes are mutated. Figure 6 shows that the reporter plasmid is efficiently repressed when the distance between the NrdR boxes is 15-17 bp, and also repressed at a distance of 26±1 bp. In contrast, the reporter plasmid appears derepressed when the distance between the NrdR boxes is 14 bp and shorter or between 18-24 bp. In general, our *in vivo* results (Fig. 6) are in good agreement with the *in vitro* results obtained with synthetic dsDNA fragments (Fig. 5C, Table 6).

**Figure 6.**
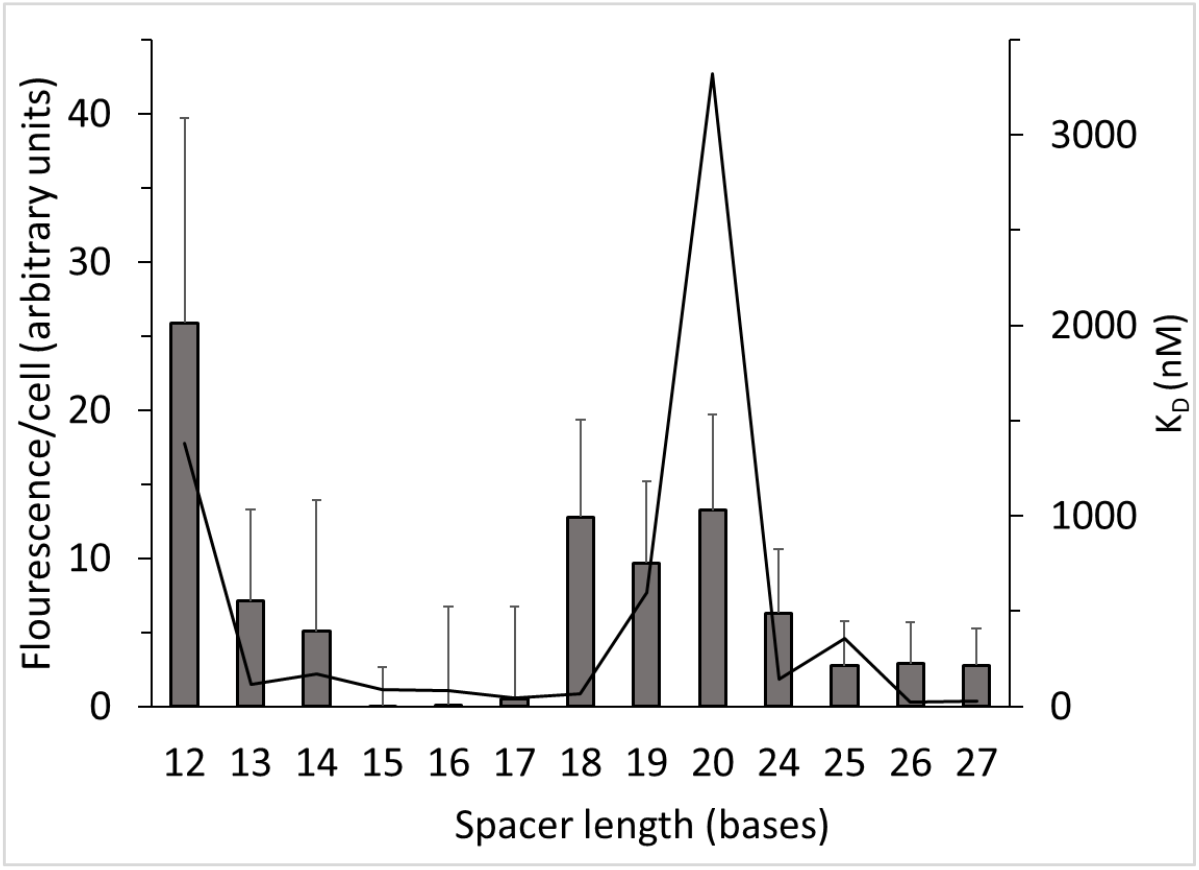
*In vivo* transcription of synthetic versions of the *E. coli* nrdHEF promoter with varying spacers between NrdR boxes, monitored by using GFP reporter gene. Standard deviations were calculated based on seven biological replicates. For comparison the K_D_ curve of *in vitro* binding of EcoNrdR to synthetic dsDNA fragments from figure 5C is included. A reporter plasmid with two mutated NrdR boxes at the wild type distance had a fluorescence/cell of 118 ±79.

## Discussion

In this report we have studied the RNR-specific transcriptional repressor NrdR from two bacterial pathogens of the phylum *Bacillota*: *L. monocytogenes* and *Streptococcus pneumoniae*. The purified NrdR proteins from *L. monocytogenes* and *Streptococcus pneumoniae* require the binding of cofactors dATP and ATP to their ATP-cones to form DNA-binding tetramers. ATP-loaded LmoNrdR and SpnNrdR were both unable to bind DNA, but formed oligomers of different sizes, similarly to ATP-cone containing RNRs, in which the active state is conserved and the inactive is not (30). These results are in agreement with previous reports of NrdR from *E. coli* and *Streptomyces coelicolor* (8,28) and suggest conservation of the NrdR mechanism throughout the bacterial domain.

A majority of NrdR binding sites upstream of RNR operons consist of two NrdR binding boxes separated by 15 to 16 bp (8,28). In addition, the *nrdAB* operon in *L. monocytogenes* has two pairs of NrdR boxes (Fig 2). LmoNrdR binds almost 5 times weaker to the *nrdAB* distal NrdR box pair separated by 15 bp compared to the *nrdAB* proximal box pair separated by 16 bp, and it binds approximately 2 times weaker to the *nrdAB* proximal box pair compared to the *nrdDG* box pair separated by 16 bp. The difference in binding to the distal and proximal box pairs may be even larger *in vivo* depending upon other factors that may bind to the upstream region of the *nrdAB* operon.

In the genome of *Streptococcus pneumoniae* the *nrdHEF* and *nrdDG* operons contain NrdR boxes separated by exceptionally long spacers (Fig. 1). The majority of Streptococci species have a distance of 25-26 bp between their NrdR boxes, but approximately a fifth, including *Streptococcus thermophilus*, have spacers of 16 bp (Table 5). Since the binding of LmoNrdR to its native promoter is orders of magnitude stronger compared to the binding of SpnNrdR to its native promoter, we asked whether the difference is due to the longer spacer regions in *Streptococcus pneumoniae*. Surprisingly, SpnNrdR binds 13 times stronger to NrdR boxes with a 16 bp spacer length compared to its native spacer length. Plausibly, the long spacer between NrdR boxes in *Streptococcus pneumoniae* is needed to relieve RNR repression, or other factors present *in vivo* may promote binding of SpnNrdR to its native binding sites.

These results triggered a general study of spacer length requirements for four different NrdR proteins to a synthetic NrdR box dsDNA fragment where the distance between the two boxes varied between 12 and 27 bp. The four NrdR proteins stem from the *Actinomycetia* (ScoNrdR), *Bacilli* (LmoNrdR and SpnNrdR) and *Gamma Proteobacteria* (EcoNrdR). In general, the strongest binding for all four NrdR proteins was obtained with a spacer length of 15, 16 or 17 bp. A previously solved structure of ScoNrdR bound to a DNA fragment corresponding to the *nrdRJ* NrdR boxes showed a constrained DNA helix with two sharp 90° kinks each covering one of the boxes to which two monomers of the tetrameric NrdR bound via their Zn-ribbons (8). We were hence expecting a narrow spacer length requirement, but surprisingly all four NrdR proteins tested tolerated a spacer length of 3-4 bp around its preferred box distance (Fig. 5). Beyond that the binding strength decreased rapidly and at its extreme was 300 times weaker compared to the strongest binding. Interestingly, all proteins had another K_D_ minimum, i.e. better binding, at approximately 24-26 bp spacer length (Fig. 5), which would correspond to one additional turn of the dsDNA helix compared to the optimal shorter spacer length. The weakest binder for all proteins appeared to be a spacer length of 20 bp corresponding to an addition of a half turn of a dsDNA helix. These results suggest that the optimal distance between the center of the two boxes in the binding motif is an integer of dsDNA turns, in the common case 3 turns, but 4 turns are also tolerated.

When supplementing our *in vitro* results with *in vivo* studies using a reporter containing the *E. coli nrdEF* promoter we observed a general consistency. The reporter was strongly derepressed at spacer lengths of 15-17 bp and also markedly at spacer lengths 25-27 bp. The least repressed spacer lengths were 12 bp and 18-20 bp, but the operon exhibited some degree of repression across all tested spacer lengths. Gene expression reached a maximum of 22% for a 12 bp spacer length compared to a mutant, in which the NrdR boxes were mutated. This observation aligns with NrdR exhibiting low affinity for DNA fragments with unfavorable spacer lengths but failing to bind to DNA with mutated NrdR boxes (8). Plausibly, this residual affinity to non-optimal binding sites aids NrdR to search DNA for its cognate binding sites. The presence of the Fur repressor in close proximity to this operon’s binding site may also contribute to repression *in vivo*.

Few transcription factors have been reported to recognize binding sites separated by variable spacer lengths (31-34), but the distance requirements between promoter elements is particularly well studied in sigma factors, where the efficiency of RNA polymerase (RNAP) binding to promoters depends on both the sequence of the binding sites and the spacer between them (35,36). Because DNA is helical, relative orientations of the −35*/*−10 binding sites are different in spacers of different lengths and can change in presence of torsional stress, modulating RNAP activity. Moreover, DNA supercoiling varies throughout the bacterial growth cycle as well as in response to a variety of physiological conditions (37), therefore the spacer acts as an environmental sensor (38,39). Bacteria often cope with a frequently changing environment, and many encode more than one RNR gene (40) with NrdR boxes in their promoter regions that may vary in sequence, spacer length and position relative to promoter elements (Fig. 1 and 2). Hypothetically, NrdR could sense the environmental conditions via the DNA supercoiling state in each RNR promoter and bind differentially to NrdR boxes in response to conditions reflecting a required stringency of RNR regulation. DNA supercoiling and likely also NrdR binding are affected by the presence of RNAP and other regulatory proteins (37,41). NrdR binding and the degree of repression will therefore depend on all the above making RNR regulation a fine-tuned interplay between NrdR, RNAP and other transcription factors binding in the region.

We recently showed that the relative orientation of ATP-cone domains is significantly different in *Streptomyces coelicolor* and *E. coli* NrdRs bound to DNA (28). We speculate that non-conserved sequences found in the linker region between the zinc finger and ATP-cone domains and/or at the C-terminal ends (Supplementary Fig. 7) determine how flexible an NrdR protein is towards the spacing between NrdR boxes. Notably, flexible interdomain linkers were shown to effectively uncouple DNA-binding domains from constraints imposed by dimerization in different members of the lacI family of regulators (42-44) and a study of zinc-finger nucleases established a direct relationship between the accepted spacing between its DNA binding sites and a short interdomain linker between the DNA-binding and nuclease domains (45).

The ATP-cone is almost exclusively found in all NrdR repressor proteins and in 60% of all RNR proteins (8,46). In NrdRs the Zn-finger domain is connected to the ATP-cone via a short linker region, and in RNRs the N-terminal ATP-cone is connected to the rest of the protein via a linker. Whereas NrdRs have to bind both ATP and dATP to form the DNA-binding tetramer, ATP-cones in RNRs only bind either ATP or dATP (47). In class I RNRs, enzyme inhibition is induced by binding of dATP to the ATP-cone, which causes oligomerization involving the ATP-cone. Recently, inhibition of the class III RNR enzyme activity was shown to be mediated by disordering of a C-terminal domain and a conserved loop adjacent to the ATP-cone linker (48). The common denominator in both NrdRs and RNRs seems to be the versatile ATP-cone domains that based on their flexible connections to the rest of the proteins can orchestrate a variety of gross conformational changes and specific oligomeric states. It is interesting that the same nucleotide binding domain is utilized by both RNRs and their specific repressor NrdR to monitor the physiological balance of nucleotide pools. Our results may contribute to the development of antimicrobial agents based on NrdR and its RNR regulons, improve prediction of NrdR binding sites in bacteria, and help understanding the function of transcription factors in general.

## Data availability

The data underlying this article are available in the article and in its online supplementary material.

## Acknowledgements

The authors thank Henrietta Nielsen, Stockholm University, for letting us use the MST instrument, and Annette Roos, SciLifeLab, Uppsala University for her assistance with using the Monolith NT. Automated MST instrument, Ornella Bimaï and Melis Dogrusöz for their help with handling the NrdR protein and MST experiments.

