## supplemental tables and figures for "Strong conservation of spacer lengths in NrdR repressor DNA binding sites"

**Supplementary table 1: Strain-specific oligonucleotides used in MST experiments (underlining indicates position of NrdR boxes).**

| MST oligonucleotide | ssDNA sequence |
|---|---|
| Lmo_nrdAB_distal_sense | 5’-Cy5-CTTAACTACTAAATATAGAAACTAGCGAAAAAATAGGCACTAGAAGTAGTAGCATTC-3’ |
| Lmo_nrdAB-_distal_antisense | 5’-Cy5-GAATGCTACTACTTCTAGTGCCTATTTTTTCGCTAGTTTCTATATTTAGTAGTTAAG-3’ |
| Lmo_nrdAB_proximal_sense | 5’-Cy5-CCAGAGTACTATATCTTGCATTCACAATGAAAAAGTTATACTATATATAGTATCGAAC-3’ |
| Lmo_nrdAB_proximal_antisense | 5’-GTTCGATACTATATATAGTATAACTTTTTCATTGTGAATGCAAGATATAGTACTCTGG-3’ |
| Lmo_nrdDG_sense | 5’-Cy5-CAATAAAACTTCATCTTGTGTTATATTATTAAAGCAGACACAATATATAGTTAGTGGG-3’ |
| Lmo_nrdDG_antisense | 5’-CCCACTAACTATATATTGTGTCTGCTTTAATAATATAACACAAGATGAAGTTTTATTG-3’ |
| Spn_nrdHEF_1,2_sense | 5’-Cy5-TACAAACACTATATATTGTGTTTATATATAAAAACAACAAAAGAACACCACTATATTTAGTTCAAATC-3’ |
| Spn_nrdHEF_1,2_antisense | 5’-GATTTGAACTAAATATAGTGGTGTTCTTTTGTTGTTTTTATATATAAACACAATATATAGTGTTTGTA-3’ |
| Spn_nrdHEF_0,1_sense | 5’-Cy5-TTTAAAAACAATTTATAGGATAATCATTGCTATTTCACAATACAAACACTATATATTGTGTTTATA-3’ |
| Spn_nrdHEF_0,1_antisense | 5’-TATAAACACAATATATAGTGTTTGTATTGTGAAATAGCAATGATTATCCTATAAATTGTTTTTAAA-3’ |
| Sth_nrdHEF_sense | 5’-Cy5-TCAAAACACAAGATATAGGGTTGTGTTTCAATACTAAACACAAAATATTGTATTCCTT-3’ |
| Sth_nrdHEF_antisense | 5’-AAGGAATACAATATTTTGTGTTTAGTATTGAAACACAACCCTATATCTTGTGTTTTGA-3’ |
| Spn_nrdDG_sense | 5’-Cy5-AACCAAATCAATATCTTGTGTATTTTTAAAAATTTTAGGATTTTTAACACAAGATATTGATTTTCCT-3’ |
| Spn_nrdDG_antisense | 5’-Cy5-AGGAAAATCAATATCTTGTGTTAAAAATCCTAAAATTTTTAAAAATACACAAGATATTGATTTGGTT-3’ |
| Sth_nrdDG_sense | 5’-Cy5-TCGAAAATCAATATATAGTGGTTGAGATAAAAATATACCACAATATATTGATTTTCCT-3’ |
| Sth_nrdDG_antisense | 5’-AGGAAAATCAATATATTGTGGTATATTTTTATCTCAACCACTATATATTGATTTTCGA-3’ |
| Negative_control_sense | 5’-Cy5-GATTTGCTGAAAACCTTGTTGTAATCATTGTTTAGACACTTTTCGTAATAAAACCAG-3’ |
| Negative_control_antisense | 5’-CTGGTTTTATTACGAAAAGTGTCTAAACAATGATTACAACAAGGTTTTCAGCAAATC-3’ |

**Supplementary table 2. Oligonucleotides used for analyzing the effect of spacer length on NrdR binding.** Red font indicates the spacer between the palindromic regions of NrdR boxes; underlining indicates position of NrdR boxes. Deleted nucleotides are shown in strike through lowercase gray while the added ones are shown in green color.

| Spacer length (bp) | Sequence (sense oligo) |
|---|---|
| **12** | 5′-Cy5-GCCAAACACAACATCTAGTGGTTGGATAG~~cgt~~GAGCACACAACATCTAGTGGACCTC-3′ |
| **13** | 5′-Cy5-GCCAAACACAACATCTAGTGGTTGGATAG~~cg~~TGAGCACACAACATCTAGTGGACCTC-3′ |
| **14** | 5′-Cy5-GCCAAACACAACATCTAGTGGTTGGATAGC~~g~~TGAGCACACAACATCTAGTGGACCTC-3′ |
| **15** | 5′-Cy5-GCCAAACACAACATCTAGTGGTTGGATAGCGTGAGCACACAACATCTAGTGGACCTC-3′ |
| **16** | 5′-Cy5-GCCAAACACAACATCTAGTGGTTGGATAGCGTGAAGCACACAACATCTAGTGGACCTC-3′ |
| **17** | 5′-Cy5-GCCAAACACAACATCTAGTGGTTGGATAGCGTGAAGGCACACAACATCTAGTGGACCTC-3′ |
| **18** | 5′-Cy5-GCCAAACACAACATCTAGTGGTTGGATAGCGTGAAGTGCACACAACATCTAGTGGACCTC-3′ |
| **19** | 5′-Cy5-GCCAAACACAACATCTAGTGGTTGGATAGCGTGAAGTAGCACACAACATCTAGTGGACCTC-3′ |
| **20** | 5′-Cy5-GCCAAACACAACATCTAGTGGTTGGATAGCGTGACAGTAGCACACAACATCTAGTGGACCTC-3′ |
| **24** | 5′-Cy5-GCCAAACACAACATCTAGTGGTTGGATAGCGTGAATTCACCTAGCACACAACATCTAGTGGACCTC-3′ |
| **25** | 5′-Cy5-GCCAAACACAACATCTAGTGGTTGGATAGCGTGAAGTTCACCTAGCACACAACATCTAGTGGACCTC-3′ |
| **26** | 5′-Cy5-GCCAAACACAACATCTAGTGGTTGGATAGCGTGAAGTTCCACCTAGCACACAACATCTAGTGGACCTC-3′ |
| **27** | 5′-Cy5-GCCAAACACAACATCTAGTGGTTGGATAGCGTGAAGTTACCACCTAGCACACAACATCTAGTGGACCTC-3′ |

**Supplementary table 3. Oligonucleotides (antisense) used for analyzing the effect of spacer length on NrdR binding.**

| Spacer length (bp) | Sequence (antisense oligo) |
|---|---|
| **12** | 5′-GAGGTCCACTAGATGTTGTGTGCTCCTATCCAACCACTAGATGTTGTGTTTGGC-3′ |
| **13** | 5′-GAGGTCCACTAGATGTTGTGTGCTCACTATCCAACCACTAGATGTTGTGTTTGGC-3′ |
| **14** | 5′-GAGGTCCACTAGATGTTGTGTGCTCAGCTATCCAACCACTAGATGTTGTGTTTGGC-3′ |
| **15** | 5′-GAGGTCCACTAGATGTTGTGTGCTCACGCTATCCAACCACTAGATGTTGTGTTTGGC-3′ |
| **16** | 5′-GAGGTCCACTAGATGTTGTGTGCTTCACGCTATCCAACCACTAGATGTTGTGTTTGGC-3′ |
| **17** | 5′-GAGGTCCACTAGATGTTGTGTGCCTTCACGCTATCCAACCACTAGATGTTGTGTTTGGC-3′ |
| **18** | 5′-GAGGTCCACTAGATGTTGTGTGCACTTCACGCTATCCAACCACTAGATGTTGTGTTTGGC-3′ |
| **19** | 5′-GAGGTCCACTAGATGTTGTGTGCTACTTCACGCTATCCAACCACTAGATGTTGTGTTTGGC-3′ |
| **20** | 5′-GAGGTCCACTAGATGTTGTGTGCTACTGTCACGCTATCCAACCACTAGATGTTGTGTTTGGC-3′ |
| **24** | 5′-GAGGTCCACTAGATGTTGTGTGCTAGGTGAATTCACGCTATCCAACCACTAGATGTTGTGTTTGGC-3′ |
| **25** | 5′-GAGGTCCACTAGATGTTGTGTGCTAGGTGAACTTCACGCTATCCAACCACTAGATGTTGTGTTTGGC-3′ |
| **26** | 5′-GAGGTCCACTAGATGTTGTGTGCTAGGTGGAACTTCACGCTATCCAACCACTAGATGTTGTGTTTGGC-3′ |
| **27** | 5′-GAGGTCCACTAGATGTTGTGTGCTAGGTGGTAACTTCACGCTATCCAACCACTAGATGTTGTGTTTGGC-3′ |

**Supplementary table 4. Sequence of NrdR boxes and the spacer between them in fragments used for reporter gene analysis (underlining indicates position of NrdR boxes)**

| Spacer length (bp) | Sequence |
|---|---|
| 12 | 5′-TTGCTATATATTGTGTTGAATCTTTTTTCAACTACATCTAGTAT-3′ |
| 13 | 5′-TTGCTATATATTGTGTTTGAATCTTTTTTCAACTACATCTAGTAT-3′ |
| 14 | 5′-TTGCTATATATTGTGTGTTGAATCTTTTTTCAACTACATCTAGTAT-3′ |
| 15 (wild type) | 5′-TTGCTATATATTGTGTGGTTGAATCTTTTTTCAACTACATCTAGTAT-3′ |
| 16 | 5′-TTGCTATATATTGTGTTGGTTGAATCTTTTTTCAACTACATCTAGTAT-3′ |
| 17 | 5′-TTGCTATATATTGTGTTTGGTTGAATCTTTTTTCAACTACATCTAGTAT-3′ |
| 18 | 5′-TTGCTATATATTGTGTTGTGGTTGAATCTTTTTTCAACTACATCTAGTAT-3′ |
| 19 | 5′-TTGCTATATATTGTGTTGTTGGTTGAATCTTTTTTCAACTACATCTAGTAT-3′ |
| 20 | 5′-TTGCTATATATTGTGTTGGTTGGTTGAATCTTTTTTCAACTACATCTAGTAT-3′ |
| 24 | 5′-TTGCTATATATTGTGTTGGTTGTTTGGTTGAATCTTTTTTCAACTACATCTAGTAT-3′ |
| 25 | 5′-TTGCTATATATTGTGTTGGTTTGTTTGGTTGAATCTTTTTTCAACTACATCTAGTAT-3′ |
| 26 | 5′-TTGCTATATATTGTGTTGGTTTGATTTGGTTGAATCTTTTTTCAACTACATCTAGTAT-3′ |
| 27 | 5′-TTGCTATATATTGTGTTGGTTTTGATTTGGTTGAATCTTTTTTCAACTACATCTAGTAT-3′ |

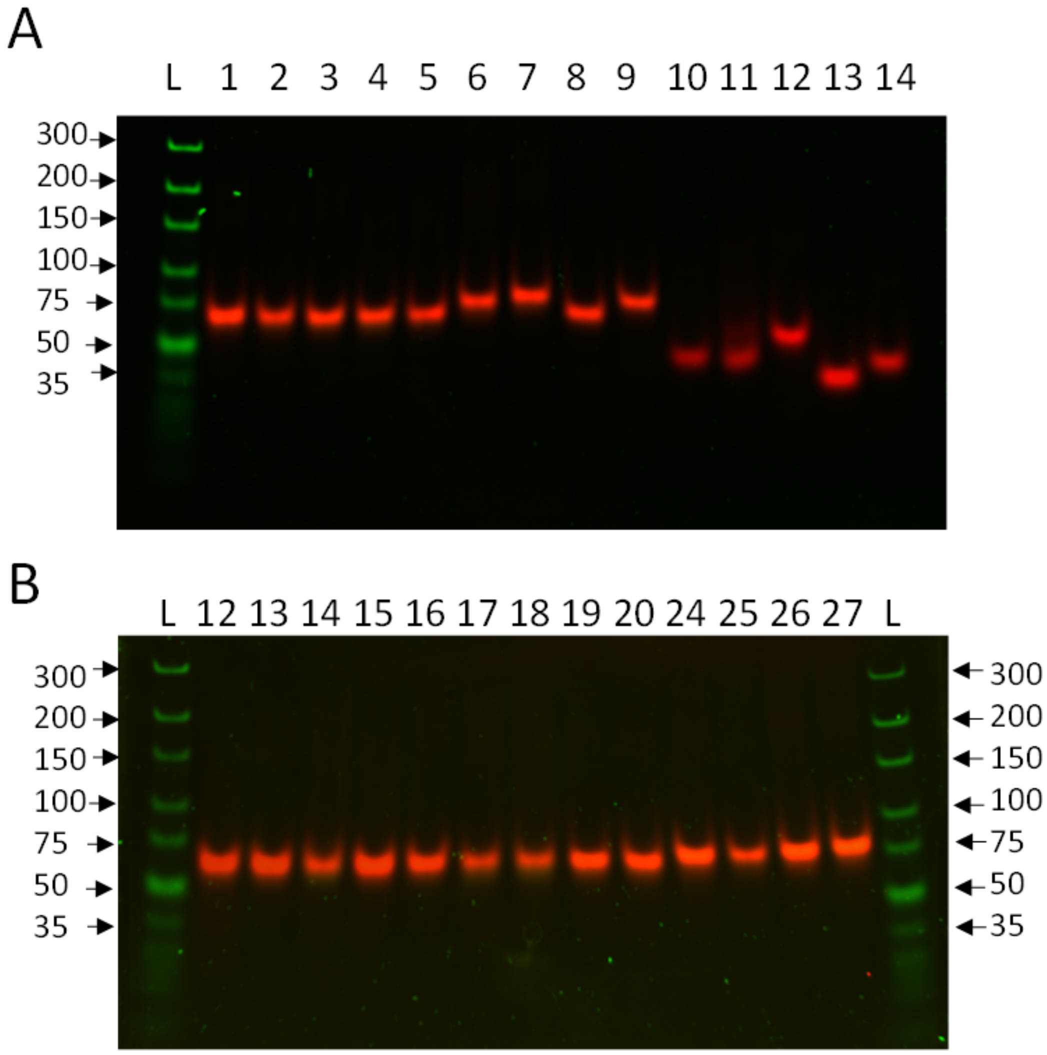


**Supplementary Figure 1: PAGE of annealed oligonucleotides used for MST. a)** L) DNA ladder; 1) LmoABdistal; 2) LmoABproximal; 3) LmoDG; 4) Negative control oligo; 5) SthAB; 6) SpnAB1,2; 7) SpnAB2,3; 8) SthDG; 9) SpnDG; 10) SthEF1,2_For; 11) SpnAB1,2_For); 12) SpnAB1,2_For; 13) SthDG_For; 14) SpnDG_For. “For” stands for single stranded sense oligo (negative control for annealing reaction). **b)** synthetic oligonucleotides with different spacer lengths: L) DNA ladder; the numbers 12-27 correspond to the length of the spacer between NrdR boxes. 1 pmol double stranded oligonucleotide was loaded on 5% polyacrylamide gel in Tris/Borate/EDTA (TBE) buffer and ran at 70 V for ~90 minutes. Gene ruler Ultra low (Thermo Fischer Scientific) was used as DNA ladder.

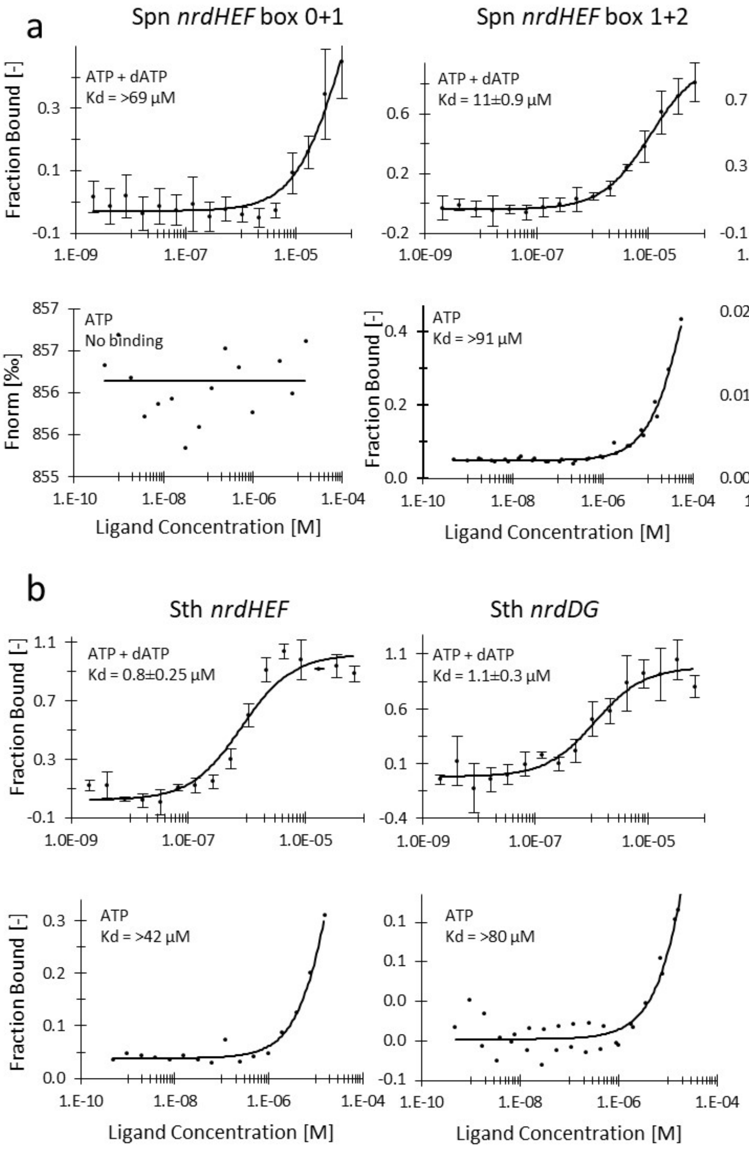


**Supplementary figure 2. Binding of *Streptococcus pneumoniae* NrdR to native *nrdHEF* and *nrdDG* promoters (a) and promoters of *Streptococcus thermophilus* (b) in the presence of different nucleotides determined by MST.** Plots of protein fraction bound vs. the concentration of ligand (NrdR) are shown. Lines represent fits of the data points using the Kd fit derived from the law of mass action. $K_D$ and standard deviations (mean values ± SD) were calculated by the analysis software using fits from at least three individual titrations. Flat lines were produced in the cases where no fit could be generated by the software and the parameters therefore were fixed as for non-binder ligands. In these traces the normalized fluorescence Fnorm (‰) from T-Jump and Thermophoresis vs. concentration of ligand are shown. In some cases the fits resulted in $K_D$s in the range of 42 - 3000 µM, but the actual $K_D$ cannot be determined, since the curves do not reach a plateau.

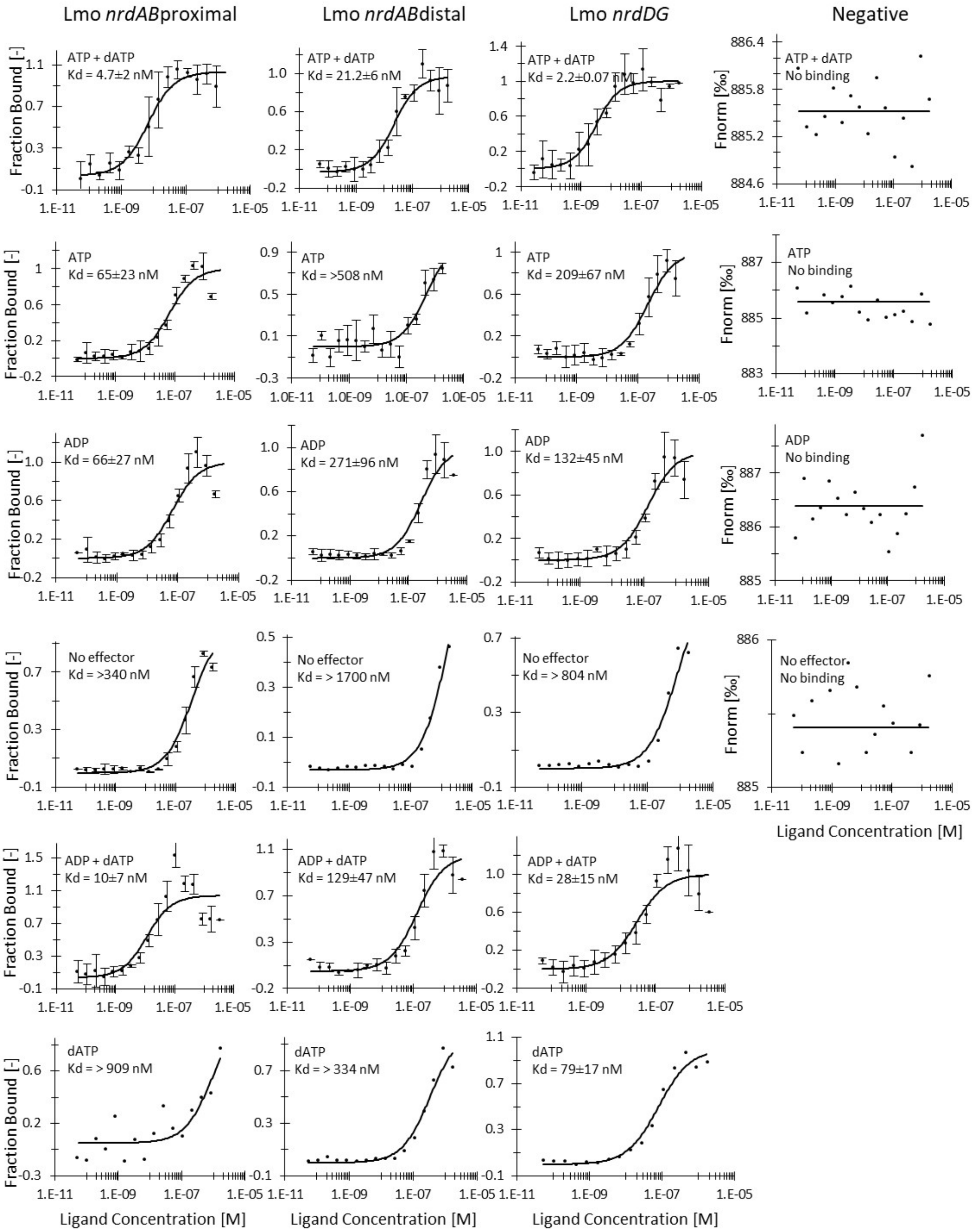
Lmo *nrdAB*proximal
Lmo *nrdAB*distal
Lmo *nrdDG*
Negative
ATP + dATP
Kd = 4.7±2 nM
ATP + dATP
Kd = 21.2±6 nM
ATP + dATP
Kd = 2.2±0.07 nM
ATP + dATP
No binding
ATP
Kd = 65±23 nM
ATP
Kd = >508 nM
ATP
Kd = 209±67 nM
ATP
No binding
ADP
Kd = 66±27 nM
ADP
Kd = 271±96 nM
ADP
Kd = 132±45 nM
ADP
No binding
No effector
Kd = >340 nM
No effector
Kd = > 1700 nM
No effector
Kd = > 804 nM
No effector
No binding
ADP + dATP
Kd = 10±7 nM
ADP + dATP
Kd = 129±47 nM
ADP + dATP
Kd = 28±15 nM
dATP
Kd = > 909 nM
dATP
Kd = > 334 nM
dATP
Kd = 79±17 nM
Fraction Bound [-]
Fnorm [‰]
Ligand Concentration [M]

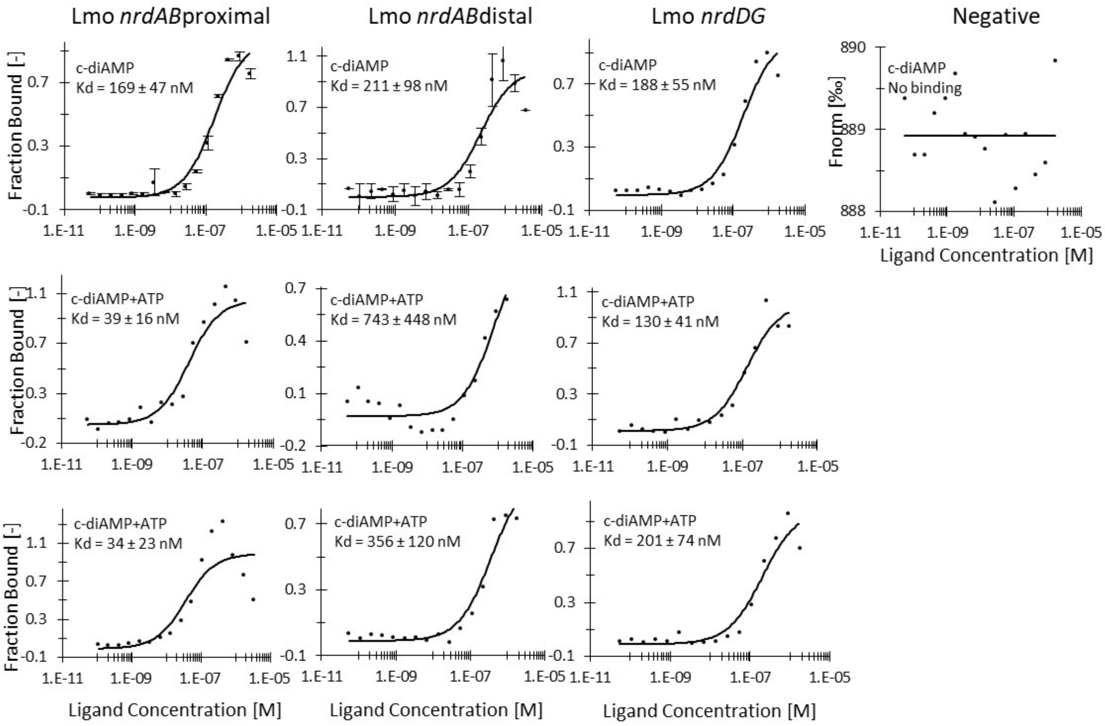


**Supplementary figure 3. Binding of *L. monocytogenes* NrdR to native promoters in the presence of different nucleotides determined by MST**. Plots of protein fraction bound vs. the concentration of ligand (NrdR) are shown. Lines represent fits of the data points using the Kd fit derived from the law of mass action. $K_D$ and standard deviations (mean values ± SD) were calculated by the analysis software using fits from at least three individual titrations. Flat lines were produced in the cases where no fit could be generated by the software and the parameters therefore were fixed as for non-binder ligands. In these traces the normalized fluorescence Fnorm (‰) from T-Jump and Thermophoresis vs. concentration of ligand are shown. In some cases the fits resulted in $K_D$s in the range of 334 - 1700 nM, but the actual $K_D$ cannot be determined, since the curves do not reach a plateau.

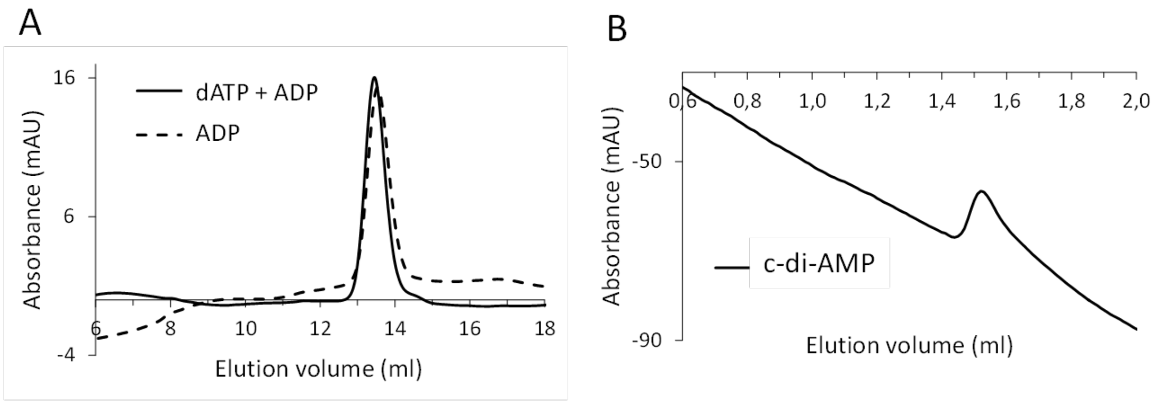


C

| | ADP | dATP +ADP | c-di-AMP |
|---|---|---|---|
| Size (kDa) | 80 | 89 | 80 |
| (# of monomers) | (4.2) | (4.7) | (4.2) |
| Oligomeric state | tetramer | tetramer | tetramer |

**Supplementary figure 4. Size exclusion chromatography of LmoNrdR in the presence of different nucleotides. a:** Chromatography performed using Superdex 200 Increase 10/300 (24 ml column) in the presence of ADP (dashed) and dATP + ADP (solid line). 100 µl of 50 µM NrdR was injected **b:** Chromatography performed on Superdex 200 Increase 3.2/300 (2.4 ml column) in presence of c-di-AMP. 25 µl of 50 µM NrdR was injected. C: summary of calculated molecular weights of LmoNrdR in presence of different effectors and its estimated oligomeric state. In parentheses is a calculated number of NrdR monomeric subunits (19 KDa).

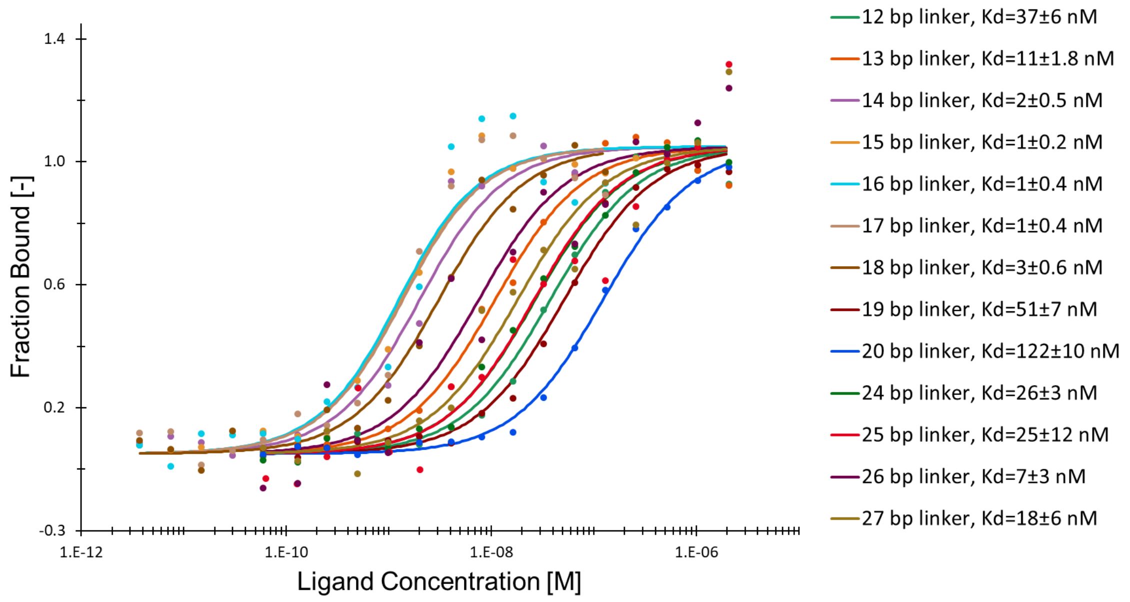


**Supplementary figure 5. Binding of LmoNrdR to synthetic NrdR boxes with varying spacer length determined by MST.** Plots of protein fraction bound vs. the concentration of ligand (NrdR) are shown. Lines represent fits of the data points using the Kd fit derived from the law of mass action. $K_D$ and standard deviations (mean values ± SD) were calculated by the analysis software using fits from three individual titrations.

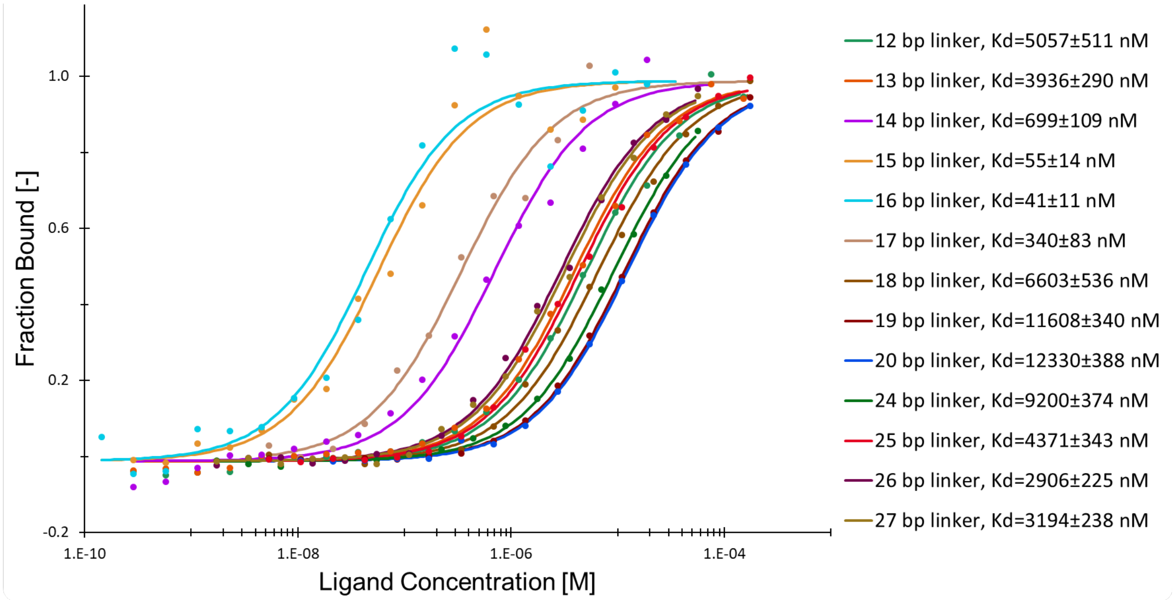


**Supplementary figure 6. Binding of SpnNrdR to synthetic NrdR boxes with varying spacer length determined by MST.** Plots of protein fraction bound vs. the concentration of ligand (NrdR) are shown. Lines represent fits of the data points using the Kd fit derived from the law of mass action. $K_D$ and standard deviations (mean values ± SD) were calculated by the analysis software using fits from three individual titrations.

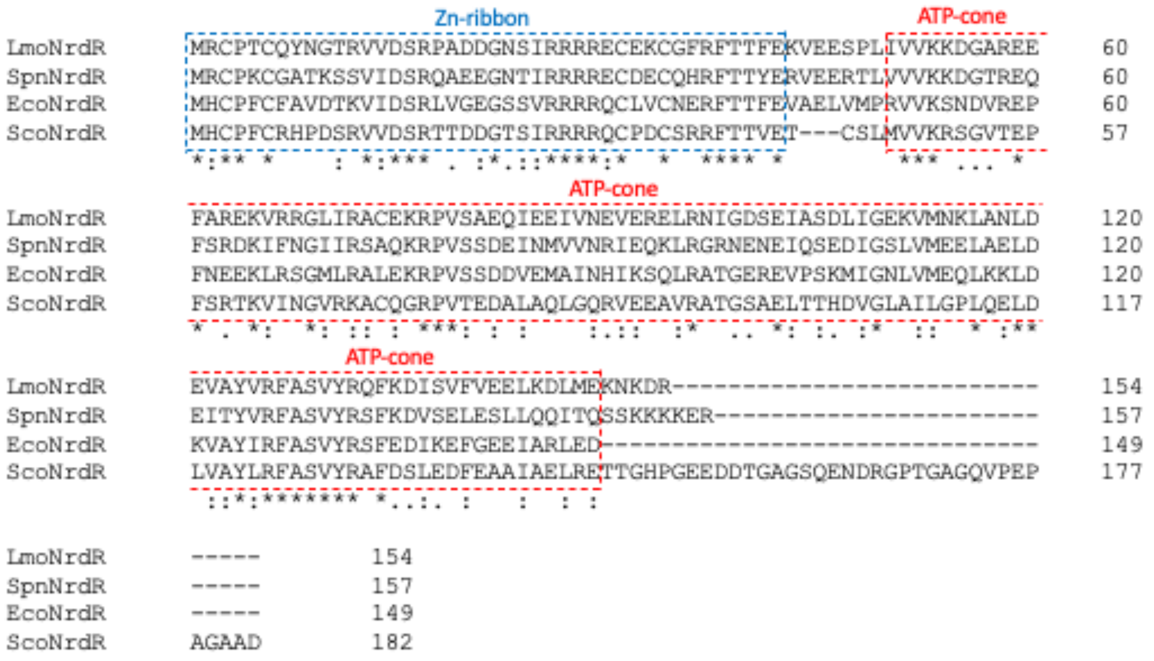


**Supplementary figure 7. Sequence alignment of NrdR proteins from *L. monocytogenes, Streptococcus pneumoniae, E. coli* and *Streptomyces coelicolor.***

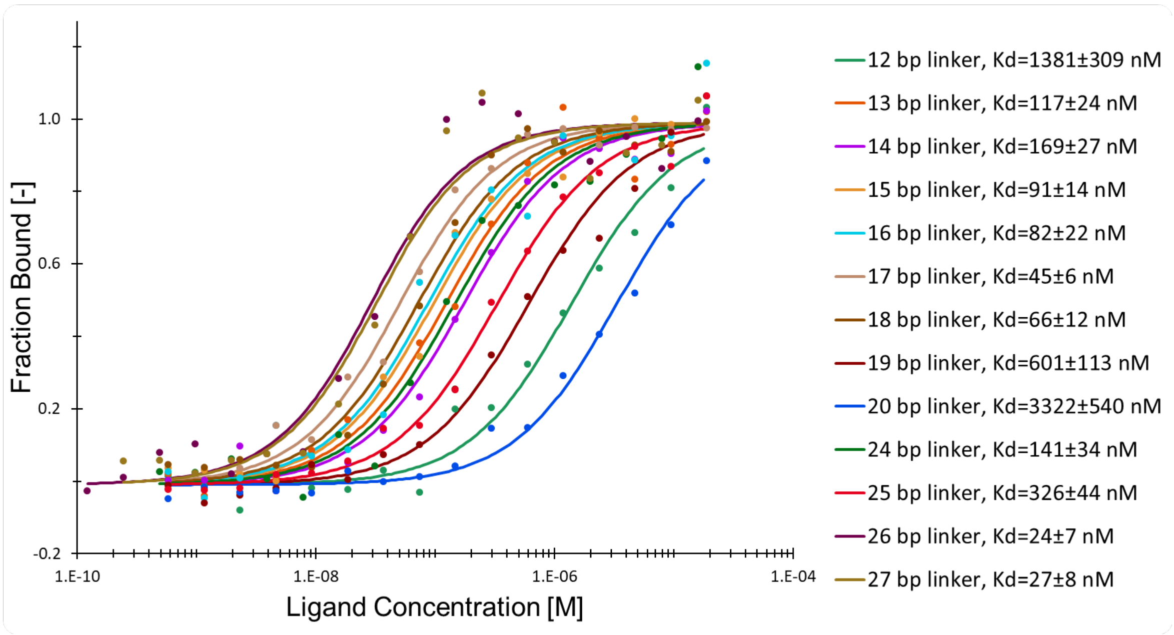


**Supplementary figure 8. Binding of EcoNrdR to synthetic NrdR boxes with varying spacer length determined by MST.** Plots of protein fraction bound vs. the concentration of ligand (NrdR) are shown. Lines represent fits of the data points using the Kd fit derived from the law of mass action. $K_D$ and standard deviations (mean values ± SD) were calculated by the analysis software using fits from three individual titrations.

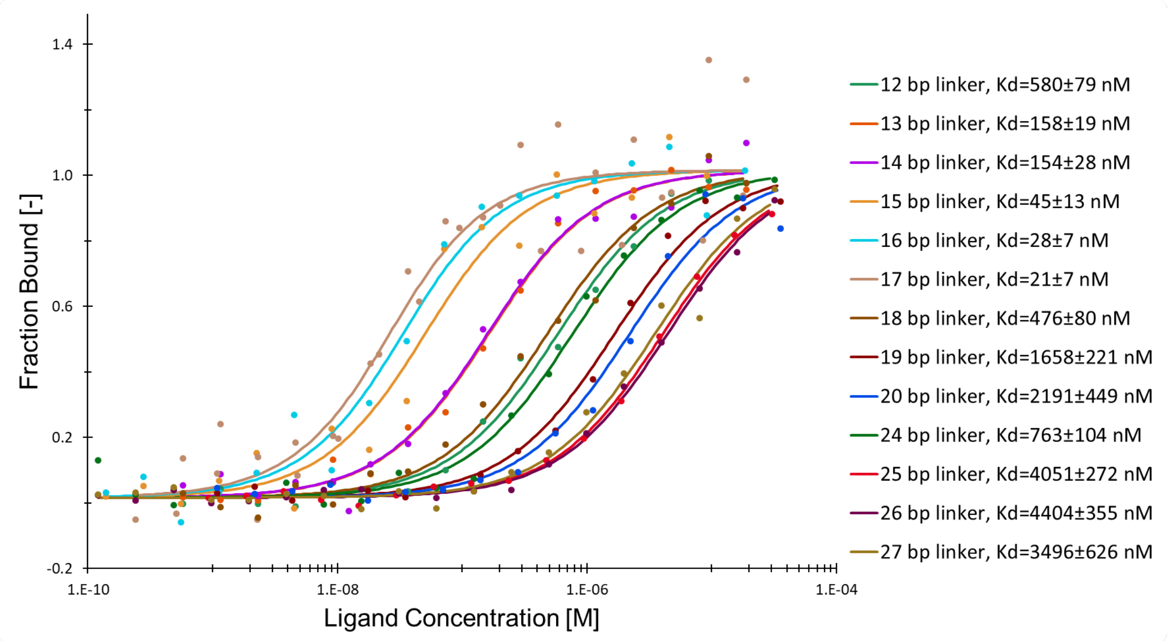


**Supplementary figure 9. Binding of ScoNrdR to synthetic NrdR boxes with varying spacer length determined by MST.** Plots of protein fraction bound vs. the concentration of ligand (NrdR) are shown. Lines represent fits of the data points using the Kd fit derived from the law of mass action. $K_D$ and standard deviations (mean values ± SD) were calculated by the analysis software using fits from three individual titrations.
